# Mixing, competence, and commitment set the stability of cell identity

**DOI:** 10.64898/2026.11.01.763472

**Authors:** Stanley E. Strawbridge, Alexander G. Fletcher

## Abstract

Cells maintain their identities in the face of molecular noise, yet can change them rapidly when cued. Here, we treat this tension as a problem in the control of a stochastic system. We represent a cell identity as a set of molecular configurations, or microstates, among which cells undergo stochastic transitions, and measure its stability by the effective rate of commitment to another identity, using a Markov model in which committed cells are removed. Mixing, competence, and commitment provide three channels for tuning this rate. They govern microstate transitions, how commitment is distributed among microstates, and the overall speed of commitment, respectively. Control over the effective commitment rate is shared between commitment and mixing in proportions that always sum to one. Interventions with the same effective commitment rate cannot be told apart by survival alone but, given the microstate transition rates, can be distinguished by the fraction of cells in each microstate or by how quickly cells sorted on permissive microstates commit. When competence varies in time, pulsed commitment signalling commits more cells than sustained signalling of equal dose, by an amount that grows with the depth of that variation, whereas with constant competence pulsing gains nothing in the models we studied. This suggests a possible function for pulsatile signalling, such as that of ERK, which the experiments we propose could test. Finally, reanalysing single-cell imaging of mouse embryonic stem cells leaving naïve pluripotency, we estimate that cells take about as long to acquire competence as they then wait to commit, and we specify experiments to identify which channel drives this exit.

## I. Introduction

Multicellular organisms depend on cells that acquire, maintain and, when necessary, change their identities. During development, identities must be established rapidly and in a coordinated way. In adult tissues, they must persist for long periods despite molecular noise. In regeneration and repair, cells must transiently regain plasticity before settling into a stable identity again. At the molecular level, a single identity comprises many fluctuating configurations of gene expression, chromatin, signalling and mechanical state. A central question in developmental and stem cell biology is how stability at the level of cell fate emerges from this dynamical complexity, and how cells can be robust to noise yet responsive to appropriate cues.

Following Waddington (Waddington, 1957), cell fate is often pictured as a landscape in which identities are attractor basins separated by barriers (Huang et al., 2005; Wang et al., 2011). Geometric approaches classify cell decisions using a small number of generic landscape geometries and the bifurcations that change them. Such models have been fitted to fate specification in the mouse blastocyst (Raju and Siggia, 2023, 2024). Non-equilibrium landscape-and-flux theories have relaxed the equilibrium assumption of the classical picture. They combine a quasi-potential that assigns an effective landscape height with a circulating probability flux. This formulation allows paths between identities, barrier heights, and first-passage times to be computed (Wang, 2015). In experiments, signals, drugs, and mechanical cues change the parameters that shape these dynamics over time. We therefore ask which parameters act as inputs and how those inputs should be scheduled to change cell identity.

To address this, we develop a framework in which a cell identity is a set of molecular configurations, or microstates, and its stability is measured by the effective rate of commitment to another identity. Mathematically, this effective commitment rate is the principal eigenvalue of a killed Markov generator, which describes microstate transition and removes cells once they commit. The mathematical ingredients of this framework are well established. Reducing a configuration space to a network of discrete states underlies Markov state models of molecular kinetics (Prinz et al., 2011; Husic and Pande, 2018). Escape from a long-lived set of states is the subject of theories of quasi-stationarity (Darroch and Seneta, 1967; Collet et al., 2013; Méléard and Villemonais, 2012) and metastability (Freidlin and Wentzell, 2012; Bovier and den Hollander, 2015). Absorbing Markov-chain methods already estimate macrostates (groups of microstates), microstates associated with commitment, and absorption probabilities (the probability that a cell eventually adopts each fate) from single-cell data (Lange et al., 2022; Weinreb et al., 2018).

Our approach is related to two existing bodies of work. The first is the control of cell fate on gene regulatory networks. There, reprogramming is cast as driving a Boolean model, in which each gene is either on or off, to a chosen attractor by fixing the states of selected nodes (Zañudo and Albert, 2015), a strategy since given a structural basis in the feedback vertex set of the regulatory graph, a set of nodes that, once fixed, determines the long-term behaviour of the network (Fiedler et al., 2013; Mochizuki et al., 2013; Zañudo et al., 2017). In that setting the dynamics are deterministic, an identity is an attractor, and control determines which attractor is reached. In ours the dynamics are stochastic, an identity is a long-lived but transient set of microstates, and the quantity being controlled is how long the current identity lasts. The two approaches are complementary, since feedback vertex sets indicate where in a network to intervene, while our framework describes what an intervention does to the lifetime of an identity and on what schedule it should be applied.

The second is the use of optimal transport and Schrödinger bridges in single-cell biology, which find the most likely way for a population of cells to move between two observed distributions of molecular configurations. These methods couple successive time points of a differentiation time course to infer ancestor–descendant relationships (Schiebinger et al., 2019; Lavenant et al., 2024), or map unperturbed onto perturbed cell populations (Bunne et al., 2023). In each case the bridge is used for inference, in that the distributions at each time are observed and the method recovers the dynamics connecting them. Here, we use the same mathematical object for design, in which the unperturbed dynamics are given, the final distribution is a target, and the solution is a schedule of interventions (Appendix F).

Our contribution is to develop the control theory of this formulation in four steps. First, we identify mixing, competence, and commitment as three channels through which stability can be tuned. They govern microstate transitions, how commitment is distributed among microstates, and the overall speed of commitment, respectively. With the competence profile fixed, the proportional sensitivities of the effective commitment rate to commitment and uniformly scaled internal transition rates sum to one (Section III). Second, we show that interventions acting through different channels can produce the same long-term effective commitment rate and identify the measurements needed to distinguish them (Section IV). Third, we treat the channels as time-dependent inputs. In the models studied, appropriately timed pulses outperform sustained commitment signalling of equal dose only when competence varies in time (Section V). This suggests a possible function for pulsatile signalling in developmental systems. Finally, we apply the framework to mouse embryonic stem cells leaving naïve pluripotency to estimate the relative times spent acquiring competence and waiting to commit. We propose experiments to identify which channels control this particular commitment process (Section VI).

## II. A killed Markov chain links microstate dynamics to identity stability

In this section, we set out the description of a cell identity on which the rest of the paper builds. A cell moves among many molecular configurations that all count as the same identity, and can commit to another identity only from some of them. Table 1 summarises the terms we use and what each means for a population of cells.

**Table 1.**
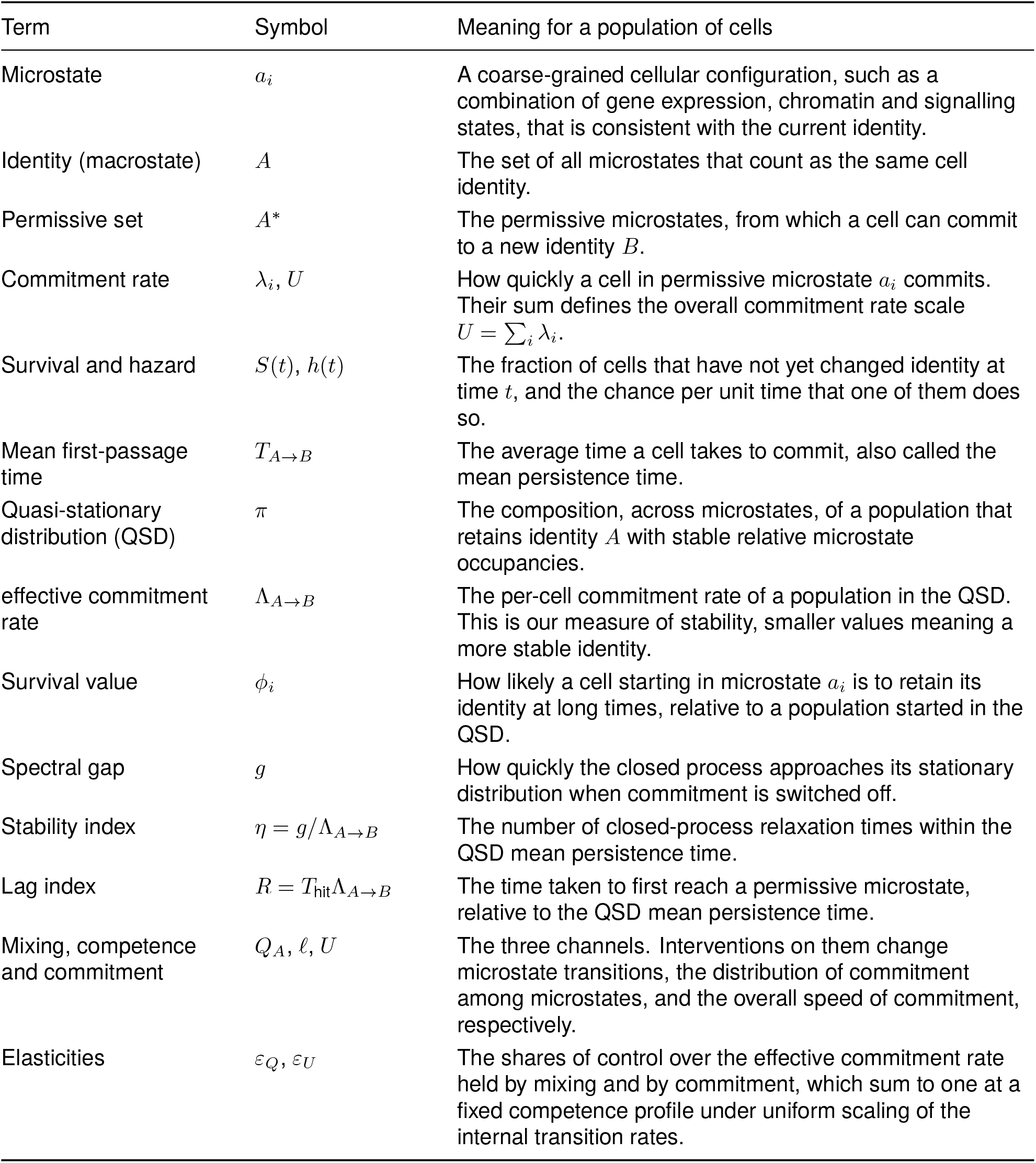
Key terms. The symbol for each term and what it means for a population of cells.

| Term | Symbol | Meaning for a population of cells |
|---|---|---|
| Microstate | a<sub>i</sub> | A coarse-grained cellular configuration, such as a combination of gene expression, chromatin and signalling states, that is consistent with the current identity. |
| Identity (macrostate) | A | The set of all microstates that count as the same cell identity. |
| Permissive set | A<sup>*</sup> | The permissive microstates, from which a cell can commit to a new identity B. |
| Commitment rate | λ<sub>i</sub>, U | How quickly a cell in permissive microstate a<sub>i</sub> commits. Their sum defines the overall commitment rate scale U = ∑<sub>i</sub> λ<sub>i</sub>. |
| Survival and hazard | S(t), h(t) | The fraction of cells that have not yet changed identity at time t, and the chance per unit time that one of them does so. |
| Mean first-passage time | T<sub>A→B</sub> | The average time a cell takes to commit, also called the mean persistence time. |
| Quasi-stationary distribution (QSD) | π | The composition, across microstates, of a population that retains identity A with stable relative microstate occupancies. |
| effective commitment rate | Λ<sub>A→B</sub> | The per-cell commitment rate of a population in the QSD. This is our measure of stability, smaller values meaning a more stable identity. |
| Survival value | ϕ<sub>i</sub> | How likely a cell starting in microstate a<sub>i</sub> is to retain its identity at long times, relative to a population started in the QSD. |
| Spectral gap | g | How quickly the closed process approaches its stationary distribution when commitment is switched off. |
| Stability index | η = g/Λ<sub>A→B</sub> | The number of closed-process relaxation times within the QSD mean persistence time. |
| Lag index | R = T<sub>hit</sub>Λ<sub>A→B</sub> | The time taken to first reach a permissive microstate, relative to the QSD mean persistence time. |
| Mixing, competence and commitment | Q<sub>A</sub>, ℓ, U | The three channels. Interventions on them change microstate transitions, the distribution of commitment among microstates, and the overall speed of commitment, respectively. |
| Elasticities | ε<sub>Q</sub>, ε<sub>U</sub> | The shares of control over the effective commitment rate held by mixing and by commitment, which sum to one at a fixed competence profile under uniform scaling of the internal transition rates. |

### II.A. Cells retain their identity during microstate transitions and commit only from a permissive set

We represent a cell identity as a macrostate *A* = *{a*_1_, …, *a*_*n*_*}* comprising a finite set of microstates. Each microstate is a coarse-grained cellular configuration, characterised by features such as molecular composition, chromatin organisation, mechanical state and signalling context. Configurations belong to the same macrostate when they satisfy the functional criteria that define the identity. Cells move stochastically among these configurations through intrinsic fluctuations while retaining their identity (Fig. 1A). A similar distinction between an identity (a macrostate) and its constituent configurations (microstates) has been used to interpret cell fate decisions and stem cell differentiation (Moris et al., 2016; Stumpf et al., 2017). We assume that commitment to a competing identity *B* is possible only from a subset *A*^*\**^ *⊆ A*, which we call the *permissive set*. Cells in *A*^*\**^ retain their identity and may return to non-permissive configurations before committing. We call entry into the permissive set acquisition of competence and return to non-permissive microstates loss of competence. The permissive set is closely related to the transition state of Moris et al. (2016), a heterogeneous and reversible pool of cells from which cells switch fate probabilistically, but here we treat the permissive configurations as part of the original identity. The case *A*^*\**^ = *A*, in which commitment is possible from every microstate, is allowed.

**Figure 1.**
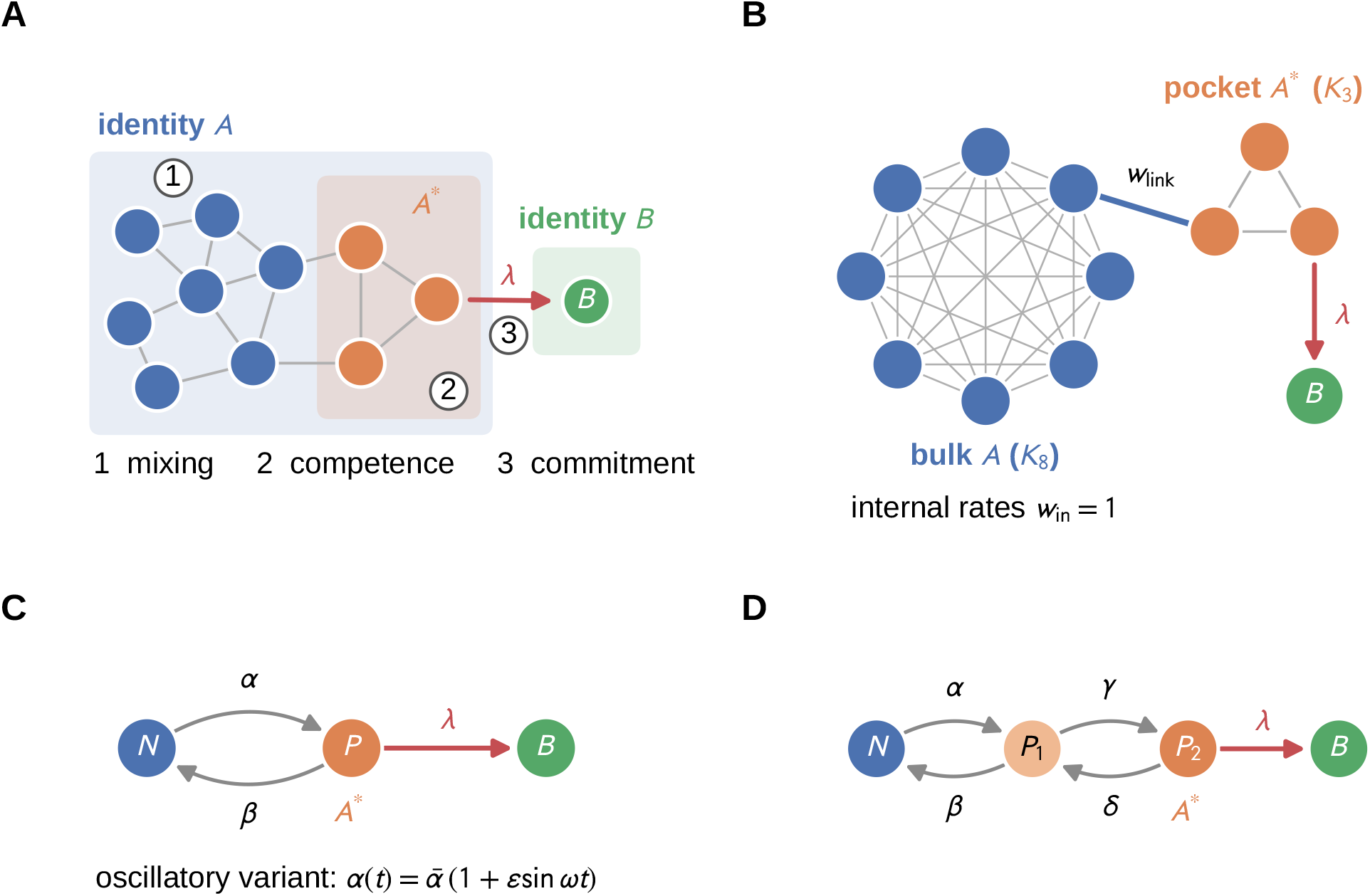
Conceptual overview of the state-space framework and the simulated network architectures. (A) A cell identity is a macrostate *A* of fluctuating microstates explored by intrinsic noise (generator *Q*_*A*_). Fate change is gated by a permissive set *A*^*\**^, from which the cell commits irreversibly (rate *λ*) to a competing identity *B*. Stability is set by three separately addressable channels, namely mixing, competence (which microstates permit commitment, and in what proportion), and commitment kinetics. (B) The 11-state test network, comprising an eight-microstate bulk clique (identity *A*) joined by a single variable-rate edge *w*_link_ to a three-microstate permissive pocket (*A*^*\**^), isolating pathway accessibility from transition-set size. (C) The running two-state model (*N ⇌ P → B*), used to compare sustained against pulsed, time-varying input. (D) The three-state line (*N ⇌ P*_1_ *⇌ P*_2_ *→ B*), used for minimum-effort (Schrödinger-bridge) steering through an intermediate state. Throughout, blue denotes non-permissive microstates, amber the permissive set *A*^*\**^, and green the acquired identity *B*. Grey edges and paired grey arrows are reversible transitions, and red arrows are irreversible commitment.

Following established approaches to representing molecular dynamics by discrete states (Prinz et al., 2011; Husic and Pande, 2018), we describe transitions within *A* by a continuous-time Markov process with generator *Q*_*A*_. Its off-diagonal entries *q*_*ij*_ *≥* 0 are the rates of transition from *a*_*i*_ to *a*_*j*_, and its rows sum to zero. Commitment from *a*_*i*_ occurs at rate *λ*_*i*_, positive on *A*^*\**^ and zero elsewhere, and *B* is absorbing, meaning that cells which reach it stay there. The probability *p*_*i*_(*t*) of occupying *a*_*i*_ at time *t* without having committed then evolves according to

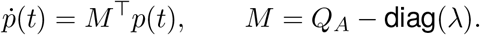

The matrix *M* generates the *killed* process, in which cells that commit are removed from the population. The survival *S*(*t*) = ∑_*i*_ *p*_*i*_(*t*) is the probability of retaining the original identity, and its hazard

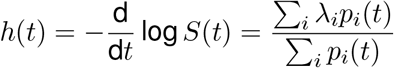

is the commitment rate averaged over the cells that remain (Appendix B.1). Put simply, *S*(*t*) is the fraction of cells that have not yet changed identity, and *h*(*t*) is the chance per unit time that one of them does so. The time at which a cell commits is its first-passage time to *B*. The mean first-passage time, which we also call the mean persistence time, 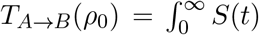 d*t*, depends on the initial distribution *ρ*_0_ of cells across microstates. In previous work, we combined parametric survival models with illustrative transition networks to examine how network organisation affects the timing of commitment (Strawbridge and Fletcher, 2026). Here, we instead express persistence directly in terms of microstate transition and commitment rates.

### II.B. The effective commitment rate measures the stability of an identity

We assume that every microstate in *A* can be reached from every other (*Q*_*A*_ is irreducible) and that *A*^*\**^ is non-empty. Among cells that have not yet committed, the distribution over microstates then converges to a unique *quasi-stationary distribution* (QSD), *π* (Darroch and Seneta, 1967). This is the distribution that remaining cells settle into once they have lost memory of where they started. It is the left principal eigenvector of *M*,

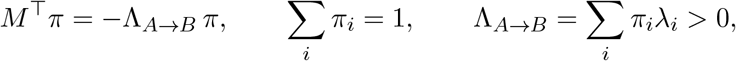

and *−*Λ_*A→B*_ is the eigenvalue of *M* with largest real part (Appendix B.2). A population that starts in the QSD keeps this distribution as cells commit, so its survival is exactly 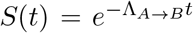 and its mean persistence time is 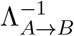 . We therefore use the effective commitment rate Λ_*A→B*_ as our measure of stability, with smaller values corresponding to a more stable identity. Populations that start elsewhere decay at the same rate in the long run, but their early hazards and mean persistence times differ. Because commitment depletes permissive configurations preferentially, *π* generally differs from the stationary distribution *π*^0^ of the closed process. With commitment switched off, this distribution satisfies 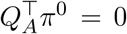. The fraction of surviving cells in permissive configurations, 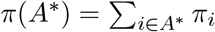, therefore depends on both the transitions within *A* and on commitment, and not only on the number of permissive microstates. In biological terms, *π* describes the composition of a population retaining identity *A* whose relative microstate occupancies have stabilised, and Λ_*A→B*_ is the effective commitment rate of that population.

### II.C. Accessibility and commitment kinetics can each limit the effective commitment rate

A model with only two configurations already captures the central trade-off. A cell must first acquire competence and then commit, and either step can limit how quickly it changes identity. For a stem cell population, *A* might be the undifferentiated state and *B* the first committed lineage, but the same description applies to any identity that cells can leave. The simplest example has one non-permissive configuration *N* and one permissive configuration *P*,

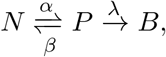

where *α* is the rate of entry into *P, β* the rate of return to *N*, and *λ* the commitment rate (Fig. 1C). The effective commitment rate is

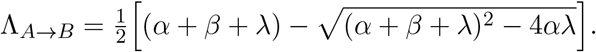

When commitment is slow compared with exchange between the two configurations (*λ « α* + *β*), the QSD approaches *π*^0^ and 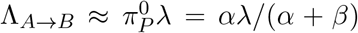. In this regime the effective permissiveness of *A* is the long-run fraction of time spent in *P*, and so depends on how readily *P* is reached and how long it is retained. As *λ* increases, Λ_*A→B*_ approaches *α*, and further increases in *λ* have a diminishing effect because access to *P* becomes limiting. Starting from *π*^0^, the mean first-passage time is *T*_*A→B*_ = (*α* + *β*)/(*αλ*) + *β*/[*α*(*α* + *β*)] (Appendix B.3). The first term is the inverse of *αλ*/(*α*+*β*), the leading-order effective commitment rate when commitment is slow, so that 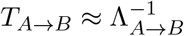 when commitment is slow relative to mixing. We also use a three-state extension, *N ⇌ P*_1_ *⇌ P*_2_ *→ B* (Fig. 1D), and an eleven-state network that separates the accessibility of the permissive set, meaning how readily cells reach it, from its size. In this network, a three-microstate permissive ‘pocket’ is joined to an eight-microstate ‘bulk’ by a single edge of rate *w*_link_ (Fig. 1B).

### II.D. Microstate mixing and access to the permissive set define distinct timescales

Whether a single rate describes the whole course of fate change depends on how quickly surviving cells lose memory of their starting configuration. For the reversible networks used here, in which there is no net circulation of cells around any loop of microstates (detailed balance), this relaxation is governed by the spectral gap *g* of *Q*_*A*_, the slowest rate at which the closed process approaches *π*^0^. Strictly, convergence of the surviving population to the QSD is governed by the spectrum of the killed generator *M*, but when commitment is slow compared with mixing the two gaps are close, and we use *g* because it depends on mixing alone. We define the stability index

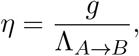

which counts the relaxation times contained in a mean persistence time. Large *η* describes a robust identity that mixes many times before cells commit, while *η ∼* 1 describes a poised identity in which commitment competes with mixing. On the eleven-state network, the hazard from a localised starting configuration relaxes to Λ_*A→B*_ on a timescale 1/*g*, and the relative error in approximating *T*_*A→B*_ by 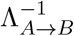 falls as 1/*η* for *η ≳* 10, for each of the three link strengths tested (Fig. S1). The error is largest near *η ≈* 6 and falls again at smaller *η*, because commitment is then limited by a single slow crossing and the effect of the starting configuration becomes small compared with the total time. Only the *η ≳* 10 branch therefore supports the single-rate description.

The spectral gap describes relaxation from a nearby configuration. A differentiation protocol, however, starts cells far from the permissive set, deep within the original identity. When the dynamics drift towards *A*^*\**^, relaxation from such a start is limited by the time taken to travel through the microstates, a timescale that the spectral gap need not reflect. We therefore also use the index

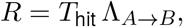

where *T*_hit_ is the mean time for the closed process to first reach *A*^*\**^ from the starting configuration. When *R «* 1, the time to first acquire competence is short compared with the mean persistence time 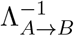, so persistence is dominated by the wait once cells are competent. When *R ≈* 1, the two are comparable, and when *R »* 1, persistence is dominated by the lag before cells become competent. In Section VI.B we show that the two indices can differ by an order of magnitude, and that only *R* can be estimated from survival data. In biological terms, *η* asks whether cells forget where they started long before they commit, and *R* asks whether the time taken to commit is spent mostly acquiring competence or mostly waiting once competent.

## III. Mixing, competence, and commitment determine identity stability

This section asks what can be changed experimentally to make an identity more or less stable, and how much each change matters.

### III.A. Accessibility changes stability even when the permissive set is fixed

The framework identifies three determinants of the stability of an identity. The first is *mixing*, the rate at which a cell explores the microstates of its identity. Mixing is set by the rates and topology of the network of transitions described by *Q*_*A*_. Highly connected regions mix rapidly, while narrow links between well-connected regions act as bottlenecks. Biologically, it may reflect chromatin organisation, transcriptional noise, cytoskeletal dynamics, metabolic state or cell–cell coupling. For example, culture conditions modulate the heterogeneity and dynamics of pluripotency-associated gene expression in embryonic stem cells (Godwin et al., 2017; Herberg et al., 2014), and local crowding reshapes cell fate through paracrine signalling (Strawbridge et al., 2025). The second is *competence*, meaning which microstates permit commitment, and in what proportion. In Waddington’s usage, competence is the ability of cells to respond to an inductive signal (Waddington, 1940), and here it refers to the configurations in which that ability is present. At quasi-stationarity, the fraction of uncommitted cells that are competent is the mass *π*(*A*^*\**^) on the permissive set. This fraction depends on microstate transitions within *A* as well as on where commitment is permitted. By competence *control* (Section III.B) we therefore mean only the redistribution of permissivity, the ability to commit, among microstates at fixed *Q*_*A*_ and total commitment rate. Interventions that change how readily *A*^*\**^ is reached alter *Q*_*A*_, and so count as mixing control. Competence may be set by chromatin priming, transcription factor availability or signalling context. Genetic screens, for instance, have identified regulators whose loss alters the capacity of embryonic stem cells to exit naïve pluripotency (Leeb et al., 2014), and signalling context can render cells refractory to differentiation cues (Mulas et al., 2017). The third is *commitment*, the speed at which cells commit from permissive microstates, given by the magnitudes of the *λ*_*i*_. Commitment corresponds to signalling strength, receptor sensitivity or pathway amplification, and are experimentally tunable. Direct reprogramming, for example, proceeds stochastically and can be accelerated without changing its eventual outcome (Hanna et al., 2009). The three determinants combine in 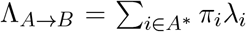. Mixing enters through *π*, competence through the support of *A*^*\**^ and the placement of permissivity within it, and commitment through *λ*.

We first asked whether the size of the permissive set alone predicts stability. On the eleven-state network, varying only the bridge rate *w*_link_ while holding the permissive set fixed changes both the competent fraction and the effective commitment rate by a factor of 38, while the size proxy |*A*^*\**^|/|*A*| stays constant (Fig. 2A). Accessibility therefore changes stability without changing the permissive set. This may explain why strong differentiation signals can fail when competence is restricted, while modest signals suffice when permissive configurations are occupied frequently. It also suggests a test. Where competence is low, increasing the accessibility of *A*^*\**^ may increase the effective commitment rate more strongly than increasing signal strength. With a fixed uniform microstate commitment rate on *A*^*\**^, the effective commitment rate is proportional to the fraction of cells occupying it.

**Figure 2.**
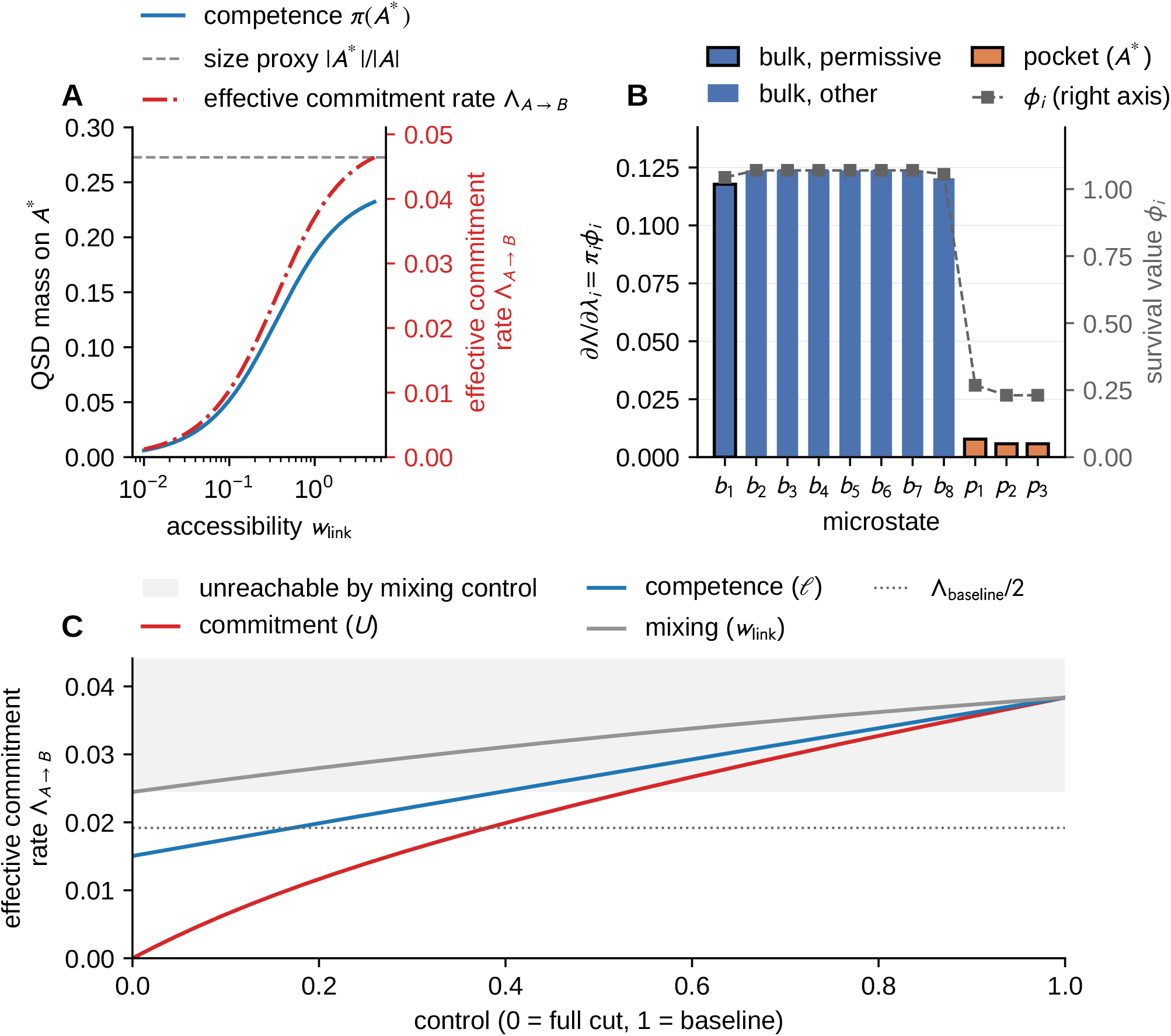
Three determinants of stability. (A) On the eleven-state network of Fig. 1B with commitment from the pocket only (*λ* = 0.2), varying only the accessibility *w*_link_ of the permissive set moves the competence *π*(*A*^*\**^) and the effective commitment rate together by a factor of 38, while the size proxy |*A*^*\**^ |/ |*A*| is constant. (B) With permissivity on the pocket and on one bulk microstate (*U* = 0.8, *w*_link_ = 0.15), the commitment sensitivity *∂*Λ_*A*_*→*_*B*_/*∂λ*_*i*_ = *π*_*i*_*ϕ*_*i*_ (bars, with permissive microstates outlined in black), which is the stationary distribution of cells conditioned never to commit, and the survival value *ϕ*_*i*_ (right axis). The permissive bulk microstate carries 86% of 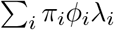, and the commitment elasticity is *ε*_*U*_ = 0.72. (C) Effective commitment rate reachable by each channel alone, from a full cut (0) to baseline (1). Mixing control has a floor at 64% of baseline, because the permissive bulk microstate stays accessible however isolated the pocket becomes.

### III.B. Occupancy and survival value determine sensitivity to intervention

To predict which intervention will most change how long cells keep their identity, we need to know which configurations matter most. Two quantities answer this question. The first is where uncommitted cells are, given by the QSD *π*. The second is how long a cell that starts in each microstate is likely to keep its identity, which we call its survival value *ϕ*_*i*_. As we show below, the product *π*_*i*_*ϕ*_*i*_ sets how sensitive the effective commitment rate is to a change in the corresponding microstate commitment rate. Mathematically, the survival value is a second eigenvector of the killed generator. Since *Q*_*A*_ is irreducible, the Perron–Frobenius theorem guarantees that the principal eigenvalue *−*Λ_*A→B*_ of *M* is real and simple, with strictly positive left and right eigenvectors. Alongside the QSD *π*, there is therefore a right eigenvector *ϕ*, satisfying *Mϕ* = *−*Λ_*A→B*_*ϕ*, which we normalise so that *π*^*⊤*^*ϕ* = 1. Whereas *π*_*i*_ is the occupancy of *a*_*i*_ among surviving cells, *ϕ*_*i*_ is its *survival value*, the relative long-run persistence of a cell that starts there, which with this normalisation is

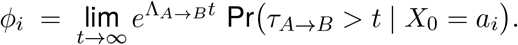

It is analogous to reproductive value in matrix population models (Caswell, 2001). To separate how strongly commitment is driven from how it is distributed among microstates, we write *λ* = *Uℓ*. Here *U* = Σ*λ*_*i*_ is the total commitment rate and *ℓ* is the competence profile, the proportions in which commitment is shared among the permissive microstates (*ℓ*_*i*_ *≥* 0, Σ_*i*_ *ℓ*_*i*_ = 1, with *ℓ*_*i*_ *>* 0 only on *A*^*\**^). A parameter point is then the triple (*Q*_*A*_, *U, ℓ*), in which mixing control varies *Q*_*A*_, commitment control varies *U*, and competence control varies *ℓ*.

#### Proposition 1

(Sensitivity decomposition). *Under the above hypotheses*, Λ_*A→B*_ *is differentiable at* (*Q*_*A*_, *U, ℓ*) *and*

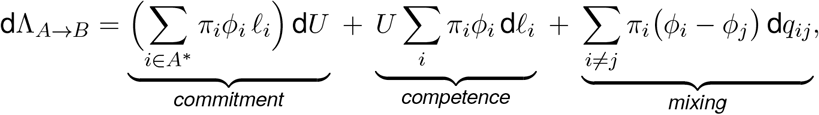

*where Σ*_*i*_ d*ℓ*_*i*_ = 0 *and the* d*q*_*ij*_ *range over off-diagonal entries of Q*_*A*_, *the diagonal adjusting to preserve zero row sums. Equivalently, in the unconstrained parameterisation*,

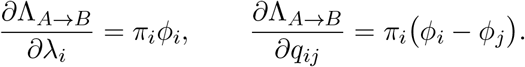

*Directional derivatives at the boundary of the simplex – that is, recruitment of a previously nonpermissive microstate into A*^*\**^ *– are one-sided but given by the same expressions*.

The proof is a first-order perturbation of a simple eigenvalue (Appendix C.1), and is the continuous-time counterpart of the sensitivity of population growth rate to the entries of a projection matrix (Caswell, 1978, 2001). Other coordinates are possible, but (*Q*_*A*_, *U, ℓ*) is a natural choice because each block corresponds to a distinct class of experimental intervention. The blocks are not orthogonal, since commitment and competence share the weight *π*_*i*_*ϕ*_*i*_. The signs of the sensitivities are also informative. Increasing any microstate commitment rate always destabilises *A*, and does so most where microstates are both frequently occupied and long-lived. The mixing sensitivity *π*_*i*_(*ϕ*_*i*_ *− ϕ*_*j*_) takes the sign of the difference in survival value along the edge, so redirecting flux from long-lived towards short-lived microstates destabilises *A*. Edges within a well-mixed region, where *ϕ* is nearly flat, have little effect even when changing them alters the mixing time substantially.

#### Corollary 1

(When the size proxy is exact). *Suppose λ*_*i*_ = *λ is uniform on A*^*\**^. *Then* 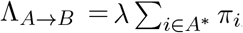, *so the competent fraction* 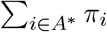 *coincides with the size proxy* |*A*^*\**^|/|*A*| *if the quasistationary distribution is uniform on A. Moreover, redistribution of permissivity within A*^*\**^ *at fixed U has no first-order effect on stability if and only if π*_*i*_*ϕ*_*i*_ *is constant on A*^*\**^.

The size proxy is therefore exact in a well-defined limit, and fails in proportion to the non-uniformity of *π*, as Fig. 2A shows when *w*_link_ is varied. In practice, counting the configurations that permit commitment is a reliable guide to stability only when cells spend roughly equal time in each configuration.

### III.C.Stability is limited by the gain of competence and commitment rate

Sensitivities to quantities with different units cannot be compared directly. We therefore use elasticities, *∂* log Λ_*A→B*_/*∂* log *θ*, which give the local proportional change in the effective commitment rate per proportional change in a parameter *θ*. For example, an elasticity of 0.5 means that a 10% increase in the parameter increases the effective commitment rate by about 5%.

#### Corollary 2

Shared control of the effective commitment rate). *Let ε*_*U*_ = (*U*/Λ_*A→B*_ )*∂*Λ_*A→B*_/*∂U be the elasticity to the total commitment rate and* 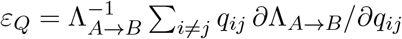 *the elasticity to a uniform rescaling of Q*_*A*_ . *Then*

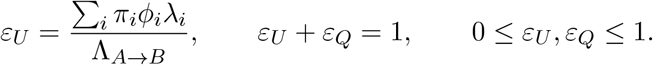

The sum rule has a simple interpretation. Speeding up every rate by 10%, both microstate transitions and commitment, increases the effective commitment rate by exactly 10%, so the effects of speeding up the two processes separately must add up to 10%. Each share lies between zero and one because speeding up either process never reduces the effective commitment rate, and never speeds it up by more than the same proportion. Neither result requires reversibility (Appendix C.4). Every identity therefore lies on a single scale. When *ε*_*U*_ is close to one, stability is limited by commitment, and changing commitment rates is the most effective way to alter it. When *ε*_*Q*_ is close to one, the effective commitment rate responds mainly to uniform scaling of the microstate transition rates. In the two-state model, the position on this scale is set chiefly by *λ*/(*α* + *β*), and only weakly by *α*/*β*. For example, *ε*_*U*_ = 0.99 when *λ* = 0.025(*α* + *β*) and 0.02 when *λ* = 25(*α* + *β*). These values recover the two limits identified in Section II.C, in which Λ_*A→B*_ is either proportional to *λ* or saturates at *α*.

The commitment weights also have a simple interpretation. With our normalisation *π*^*⊤*^*ϕ* = 1, the weights *π*_*i*_*ϕ*_*i*_ sum to one and form the stationary distribution of the *Q-process*, the dynamics of cells conditioned never to commit (Collet et al., 2013; Méléard and Villemonais, 2012). The sensitivity of the effective commitment rate to the microstate commitment rate at *a*_*i*_ is therefore the long-run fraction of time that such a cell spends in *a*_*i*_. In other words, the microstates most worth targeting are those in which long-term survivors reside. In the configuration of Fig. 2B, a single permissive microstate in the bulk accounts for 86% of Σ_*i*_ *π*_*i*_*ϕ*_*i*_*λ*_*i*_, even though three permissive microstates sit in the pocket (*ε*_*U*_ = 0.72 in this configuration). Occupancy alone does not rank microstates by their influence, because the pocket microstates are visited mainly by cells that are about to commit.

### III.D. The limits of stabilisation depend on the intervention

We next asked how far the interventions considered in each channel could change the effective commitment rate. In the same configuration, commitment and competence control can each halve the effective commitment rate, but mixing control cannot. As *w*_link_ *→* 0, the effective commitment rate along this intervention path approaches 64% of its baseline value, because the permissive microstate in the bulk remains accessible however isolated the pocket becomes (Fig. 2C). Along the competence-redistribution path considered here, with *U* fixed, the smallest effective commitment rate reached is 39% of baseline. Neither limit can be inferred from the baseline effective commitment rate alone. Each kind of intervention therefore has its own limit on how far it can stabilise an identity, so a protocol that has saturated one channel may still be improved by acting on another.

## IV. Equal effective commitment rates can conceal different control mechanisms

Different interventions can produce the same effective commitment rate. We ask when such interventions can nonetheless be told apart, and by which measurements.

### IV.A. Survival alone cannot distinguish interventions with equal effective commitment rates

Mechanistically distinct perturbations, such as supplying a transcription factor, removing a competing repressor, or altering signalling or mechanics, may produce the same effective commitment rate. Identifying such classes of interventions matters for translation, since it could allow an inefficient or cytotoxic protocol to be replaced by a safer one with the same effect on stability. Let Θ denote the set of admissible parameter points *θ* = (*Q*_*A*_, *U, ℓ*), with *Q*_*A*_ of fixed network topology.

#### Definition 1

Two parameter points are *stability-equivalent* if they have the same effective commitment rate. The *stability class* of *θ* is 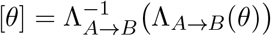.

#### Proposition 2

(Size of a stability class). *∇*Λ_*A→B*_ *≠*0 *everywhere on* Θ. *Consequently every stability class is a smooth embedded hypersurface of codimension one in* Θ, *with normal given by Proposition 1. No measurement of the asymptotic survival curve, of* Λ_*A→B*_, *or of the mean first-passage time from quasi-stationarity can distinguish two points of the same class*.

That many parameter points share an effective commitment rate is unsurprising. The useful ontent of Proposition 2 is that this degeneracy has exactly one dimension fewer than the parameter space, and that its normal direction can be computed (Appendix C.5). This is a control-theoretic form of the “sloppiness” of maps from parameters to observables, in which many combinations of parameters produce the same observable behaviour (Transtrum et al., 2015). Interventions on chromatin, signalling or mechanics that produce equal effective commitment rates are interchangeable as far as identity stability is concerned.

### IV.B. Relaxation times separate mixing control from the other channels

Alongside the effective commitment rate, we consider how quickly a perturbed population settles back to its typical composition. We ask whether this relaxation time distinguishes the channels.

#### Proposition 3

(What the spectral gap does and does not resolve). *The spectral gap g is a functional of Q*_*A*_ *alone. Hence the following hold*.

i. *Commitment control (varying U at fixed Q*_*A*_, *ℓ) and competence control (varying ℓ at fixed Q*_*A*_, *U ) both leave g invariant*.
ii. *The pair (*Λ_*A→B*_, *g) therefore cannot separate competence control from commitment control, for any perturbation of either kind*.
iii. *Generic mixing control changes g, so (*Λ_*A→B*_,*) g separates mixing control from the other two channels and cuts the stability class to codimension two*.

Proposition 3 concerns the closed generator *Q*_*A*_. The killed generator *M* = *Q*_*A*_ *−* diag(*λ*) is altered by both commitment and competence control. Its subdominant spectrum, which sets how quickly surviving cells relax onto the QSD, can therefore distinguish them. In practice, how quickly a population re-equilibrates can therefore reveal whether an intervention rewired the transitions among configurations, but not whether it changed where or how fast cells commit. Two further observables resolve the degeneracy completely.

### IV.C. Known transitions and microstate measurements can identify the control channel

If the transitions among configurations are known, either of two measurements suffices to identify the channel on which an intervention acts, namely how cells are distributed among configurations, or how long cells placed in a given configuration take to commit.

#### Proposition 4

(Identifiability from the resolved quasi-stationary distribution). *Let Q*_*A*_ *be known and let π be the resolved quasi-stationary distribution. Then the vector of microstate commitment rates satisfies*

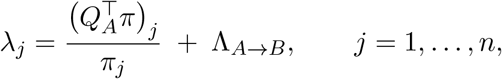

*so λ is determined by* (*Q*_*A*_, *π*) *up to a single uniform additive shift. If A*^*\**^ *≠ A, so that* 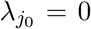 *for at least one known non-permissive microstate* 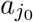, *the shift is fixed*, 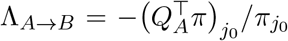, *and λ – hence the pair* (*U, ℓ*), *hence the control channel – is uniquely determined. Requiring the same value of* Λ_*A→B*_ *to be returned by every non-permissive microstate furnishes* |*A\A*^*\**^|*−*1 *independent consistency checks on the reconstruction*.

#### Proposition 5

(Identifiability from first passage). *Let Q*_*A*_ *be known and let T* = (*−M*)^*−*1^**1** *be the vector of mean first-passage times to B, T*_*i*_ *being the mean time to commitment of a cell started in a*_*i*_. *Then*

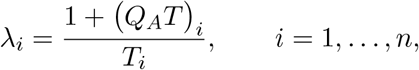

*so λ – hence* (*U, ℓ*) *and the control channel – is determined uniquely by* (*Q*_*A*_, *T* ), *with no additive ambiguity. Consequently, two parameter points with the same Q*_*A*_ *and different λ differ in the mean first-passage time from at least one microstate. Commitment and competence cuts matched in both* Λ_*A→B*_ *and g are nonetheless separated by first passage from a suitable starting state*.

Both proofs are short (Appendix C), and the two reconstructions are complementary. Proposition 4 divides by *π*_*j*_, so it is poorly conditioned for rarely occupied microstates, whereas Proposition 5 divides by *T*_*i*_, which is bounded below by the mean residence time in *a*_*i*_. The two results also call for different experiments. The first requires the resolved occupancy of an unperturbed population, whereas the second requires cells to be placed in, or sorted on, known microstates and followed until they commit. A coarse version of the second requires only a gate on the permissive set. For cells drawn from the quasi-stationary population and sorted on *A*^*\**^, the mean first-passage time satisfies 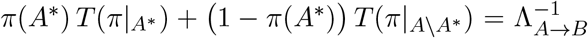. Here 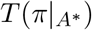 the mean time to commitment of cells sorted on *A*^*\**^, and we use the normalised contrast 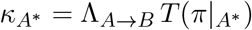, their mean time to commitment relative to that of unsorted cells, as a summary of this experiment. Both results assume that the microstate partition and at least one non-permissive microstate are known, and in practice these must themselves be estimated.

### IV.D. Sorting permissive cells can distinguish interventions with equal stability

To test these results, we gave the competence channel room to act by placing permissivity on the three pocket microstates and on one bulk microstate of the eleven-state network (*U* = 0.8, *w*_link_ = 0.15; baseline Λ_*A→B*_ = 0.0384, *g* = 0.0654). We then halved the effective commitment rate in two ways, which we refer to as cuts (Fig. 3). The commitment cut scales *U* down to 0.304 at fixed *ℓ* and *Q*_*A*_. The competence cut holds *U* and *Q*_*A*_ fixed and moves permissivity from the bulk microstate onto the pocket. Both cuts reach Λ_*A→B*_ = 0.0192 and leave *g* unchanged to machine precision (Fig. 3A), and populations started in their QSDs have identical survival curves. Their resolved occupancies nonetheless differ by a factor of 2.5 on the pocket (pocket mass 0.146 against 0.059, with a total variation distance of 0.087 between the full distributions; Fig. 3B). Since the permissive bulk microstate is occupied almost equally under both cuts, the informative gate is the pocket, and in this section and Fig. 3 the occupancy *π*(*A*^*\**^) and the sorted-start contrast 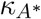 are evaluated on the pocket. Applying Propositions 4 and 5 recovers the vector of microstate commitment rates to 10^*−*14^ in both cases. The two killed generators also differ in their subdominant eigenvalue (0.132 against 0.306). Most directly, cells sorted on the pocket commit after 29.7 time units under the commitment cut, compared with 12.0 under the competence cut 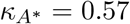 against 0.23; Fig. 3C). Under the commitment cut, cells in *A*^*\**^ wait behind a lowered gate, whereas under the competence cut, the same total commitment rate is concentrated on fewer permissive microstates, so cells that reach them commit quickly. Sorting on the bulk is uninformative (56.0 against 54.7), as expected from the near-uniform survival value there. The separation holds at every effective commitment rate that both channels can reach, and grows with the size of the cut, from a 3% difference in 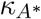 for a 2% reduction in Λ_*A→B*_ to a factor of 2.5 when the effective commitment rate is halved (Fig. 3D) . Larger cuts may therefore make these interventions easier to distinguish experimentally. On the three-state line, an intervention that reduces accessibility by lowering *α* alters *Q*_*A*_, so in our coordinates it counts as mixing control. It is then separated from a rate-matched commitment cut by the gap of *Q*_*A*_ (0.61 against 1.00), as well as by occupancy (Fig. S2C,D). In experimental terms, two interventions that delay differentiation equally can be distinguished by the fraction of cells in the pocket gate and by how quickly sorted permissive cells commit, provided that the interventions are strong.

**Figure 3.**
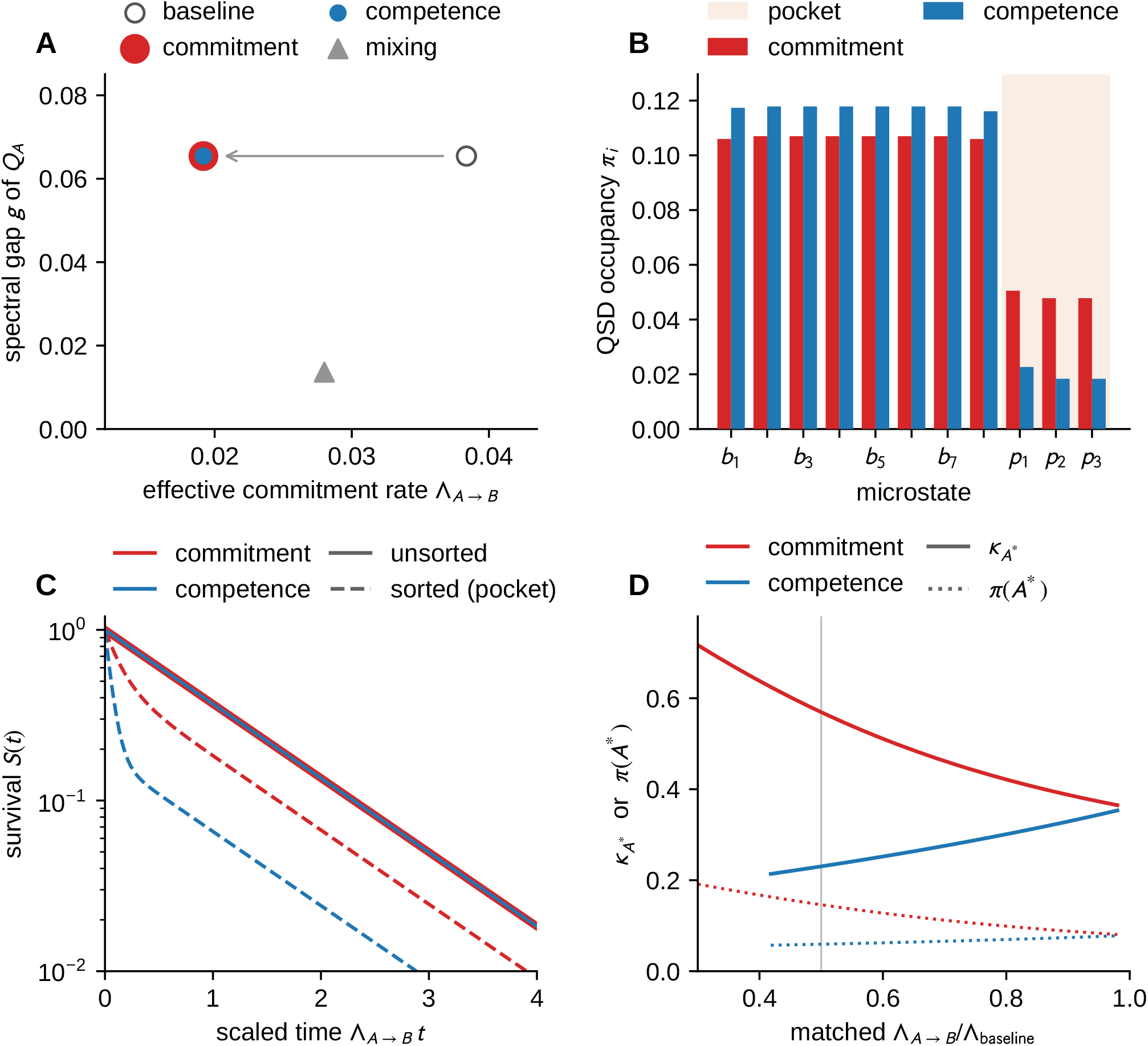
Equal effective commitment rates, different mechanisms. Configuration of Fig. 2B. **(A)** A commitment cut (scaling *U* to 0.304) and a competence cut (relocating permissivity onto the pocket at fixed *U*) both halve the effective commitment rate and leave the gap *g* of *Q*_*A*_ unchanged, so they coincide in (Λ_*A*_*→*_*B*_, *g*). A mixing cut, not rate-matched, is shown for comparison. (B) Their quasi-stationary occupancies differ by a factor of 2.5 on the pocket. (C) Survival from the QSD (solid) coincides, but survival of cells sorted on the pocket (dashed) does not, these cells committing after 29.7 against 12.0 time units. (D) Sorted-start contrast 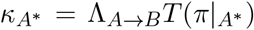 (solid) and occupancy *π*(*A*^*\**^) (dotted) for cuts matched at every value of the effective commitment rate, both evaluated on the pocket. The competence channel cannot reduce Λ_*A*_ *→* _*B*_ below 39% of baseline. Both observables separate the channels, increasingly so for deeper cuts (Propositions 4 and 5).

## V. Scheduling interventions can improve control of cell identity

We now turn from describing stability to changing it over time, as a differentiation protocol or a reprogramming cocktail does.

### V.A. Signalling inputs control microstate transitions and commitment rates

Signalling, mechanical and metabolic inputs continuously modulate the rates that determine stability. We let *u*(*t*) *∈ U ⊆* ℝ^*m*^ represent the combined effect of such inputs over a time window [0, *T* ] during which cells are responsive. The generator and microstate commitment rates then depend on the control, and the killed dynamics become

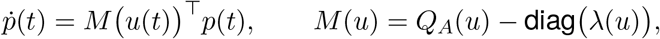

with commitment flux 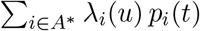. Mixing control acts on the off-diagonal entries of *Q*_*A*_(*u*), competence control on which *λ*_*i*_ may be non-zero and in what proportion, and commitment control on their magnitudes. Since *u* multiplies the state in these equations, the system is bilinear. Its reachable set, the set of microstate distributions that can be attained within the time window, has a direct biological meaning. A cell is competent when the reachable set intersects *A*^*\**^, and a competence window is an interval over which it does so.

Two further design questions are treated in Appendix F. When each rate can be tuned independently, the best protocol for either maintaining or destabilising an identity pushes every rate to one of its limits. A chosen distribution of cells can be reached with least disruption by reweighting the transitions that cells already make, which is a Schrödinger-bridge problem (Fig. S2A,B).

### V.B. Pulsed signalling can exploit variation in competence to increase commitment

Temporally structured signalling is widely reported to matter in stem cell biology. For example, a switch from symmetric to asymmetric divisions minimises the time to maturity of intestinal crypts (Itzkovitz et al., 2012), and discrete temporal control of SHH and BMP/FGF10 signalling specifies distinct tuberal hypothalamic lineages (Chinnaiya et al., 2025). Signalling activity is itself often pulsatile, as for ERK in mouse ESCs (Raina et al., 2022) and in the epiblast of the mouse blastocyst (Simon et al., 2020). None of these studies compares commitment inputs at matched integrated dose. We therefore asked a narrower question, namely when redistributing a fixed dose of commitment signal over time changes the probability of commitment.

We consider maximising the probability *J* = 1 *− S*(*T* ) of commitment by time *T*, using a commitment input *u*_*λ*_(*t*) *∈* [0, *u*_max_] that sets the microstate commitment rates, *λ*_*i*_ = *u*_*λ*_*ℓ*_*i*_, with a fixed dose 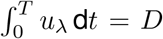. Pontryagin’s maximum principle, the standard tool for such problems, gives necessary conditions for the optimal input in terms of a costate *ψ*(*t*), with *ψ*(*T* ) = 0, where *ψ*_*i*_(*t*) is the probability that an uncommitted cell in *a*_*i*_ at time *t* commits by time *T*, and the Hamiltonian

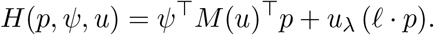

Since *H* is linear in *u*_*λ*_, the optimal input is ‘bang–bang’, switching between 0 and *u*_max_ and saturating where the switching function

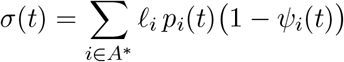

exceeds the multiplier associated with the dose constraint. The optimal input is therefore pulsatile, with pulses timed to the maxima of a costate-weighted permissive occupancy. In other words, a fixed amount of signal is best delivered at the times when many cells are both permissive and still uncommitted. Whether pulsing helps depends on whether *σ* has interior maxima. When permissive occupancy is stationary, *σ* is flat apart from end effects, and in the models we studied a constant input is then optimal. A quasi-static argument indicates why. If the input varies slowly compared with relaxation onto the QSD, then log 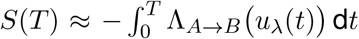, and since Λ_*A→B*_ is concave in the total commitment rate (Appendix C.4), so that each further increase in that rate gains less, Jensen’s inequality implies that, among such slowly varying inputs, a constant input of the same dose maximises *J*. A pulsing advantage therefore requires competence that varies in time, generated for example by oscillatory or transient mixing control (Appendix A discusses the scope of this result).

To quantify the effect, we used the two-state model with 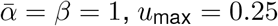, dose *D* = 1.5 and horizon *T* = 40, starting from *π*^0^. With constant mixing rates, permissive occupancy is stationary apart from a brief initial relaxation onto the QSD, *σ* has no interior maximum, and the optimal protocol gains nothing over sustained signalling (Fig. 4A,B,D). We then introduced oscillatory mixing, 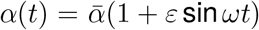 with *ω* = 0.7. Permissive occupancy now oscillates, *σ* acquires interior maxima, and the optimal input consists of pulses timed to the peaks of competence (Fig. 4C). We computed the optimum by a forward–backward sweep and report the better of that solution and the sustained protocol, so that the plotted advantage is a lower bound. The advantage grows steadily with the depth of modulation (Fig. 4D). At *ε* = 0.9 the committed fraction rises from 0.50 to 0.61, a relative gain of 23%, compared with 14% at *ε* = 0.6 and only 3.7% at *ε* = 0.2. These results suggest a possible function for pulsatile signalling. When competence varies in time, the same amount of signal commits substantially more cells if it is delivered in pulses timed to the windows of competence, and when competence is constant, pulsing confers no benefit. This has two implications for experimental design. First, cell number is unlikely to be limiting, since separating proportions of 0.50 and 0.61 requires a few hundred cells per condition and even the *ε* = 0.2 contrast requires around 10^4^, both routine by flow cytometry. Variability between biological replicates is more likely to be the limiting factor. Second, a single comparison of pulsed and sustained signalling at an unknown, possibly small, modulation depth could easily return an uninformative null result. A stronger test would vary the depth of competence modulation and ask whether the advantage rises from zero as predicted.

**Figure 4.**
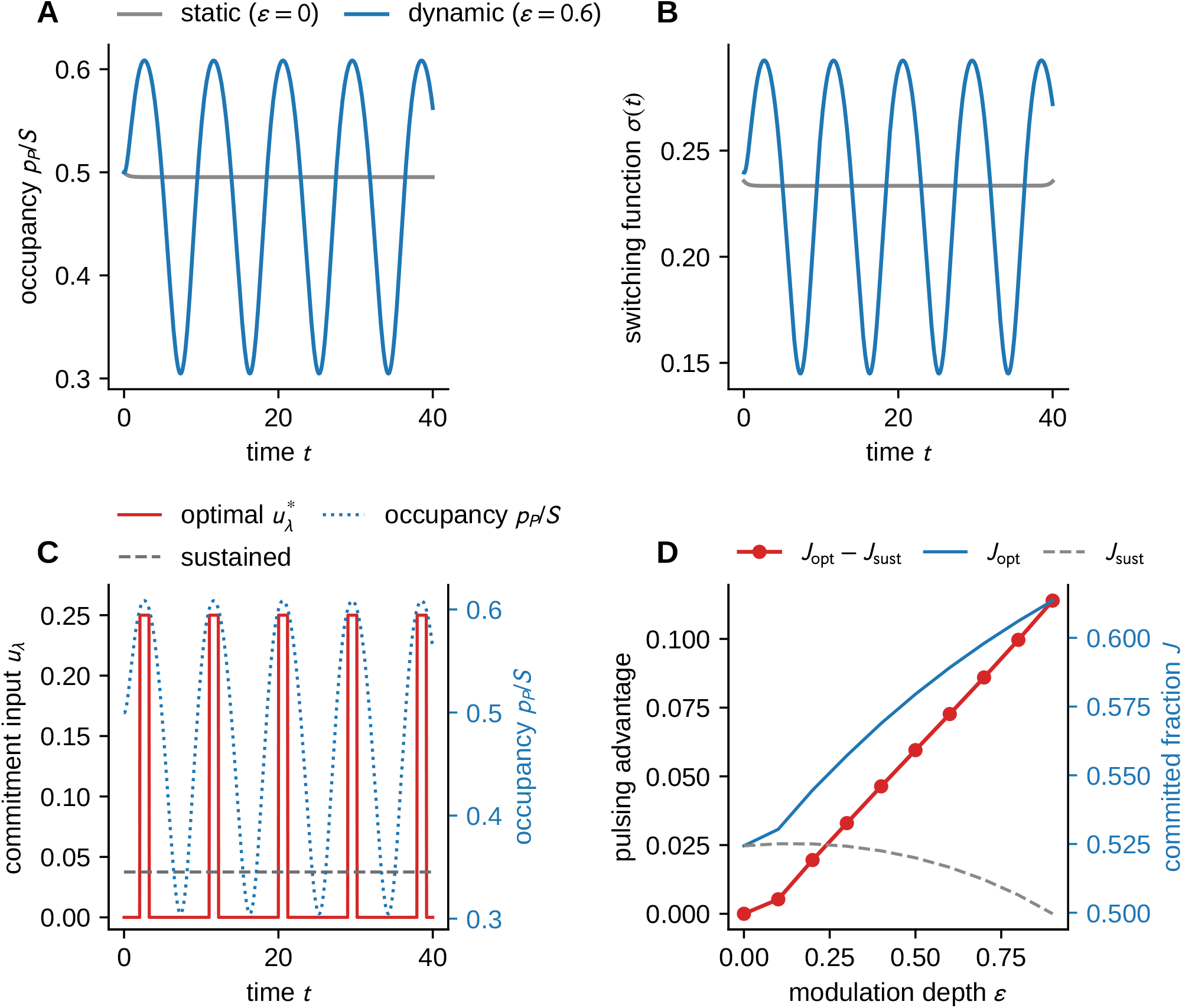
Pulsed versus sustained signalling under static and dynamic competence. Running two-state model (*N ⇌ P→B*) under a fixed integrated commitment dose. (A) Along the constant-dose reference, permissive occupancy *p*_*P*_ /*S* is stationary up to a brief initial relaxation from *π*^0^ onto the quasi-stationary distribution *π* of the killed generator with constant competence (*ε* = 0), or oscillates under time-varying mixing (dynamic competence, *ε* = 0.6). (B) The switching function *σ*(*t*), which measures the benefit of applying the commitment input at time *t*, is flat with constant competence, beyond that transient, but acquires interior maxima – “windows of opportunity” – under dynamic competence. (C) The optimal commitment input is bang–bang (solid), entrained to the competence maxima (dotted), and outperforms the sustained protocol of equal dose (dashed). (D) The pulsing advantage *J*_opt_ *− J*_sust_ is zero with constant competence, where in the models studied the sustained protocol is optimal, and grows monotonically with modulation depth *ε*.

## VI. Exit from naïve pluripotency involves a competence lag followed by commitment

We next asked where a well-characterised change in mammalian cell identity sits within this framework. We chose the exit of mouse embryonic stem cells (ESCs) from naïve pluripotency following withdrawal of the two-inhibitor (2i) ground-state condition (Kalkan et al., 2017). Exit is asynchronous between cells (Trott and Martinez Arias, 2013). Responsiveness to differentiation signals also changes around this period (Mulas et al., 2017; Semrau et al., 2017). naïve pluripotency in 2i is an in vitro state, which has been described as a developmental transition state trapped in culture (Moris et al., 2016). We therefore treat its exit as a well-characterised a model of a change in cell identity and not as a direct readout of the embryo. The macrostate (naïve identity), a permissive set and the commitment step therefore all have molecular counterparts. The destabilised *Rex1*::GFPd2 reporter provides a reporter-based measure of naïve exit through loss of GFP, allowing times to this reporter-defined endpoint to be estimated from imaging. We reanalysed long-term imaging of this reporter, using cell-level exit or censoring times together with lineage relationships (Strawbridge et al., 2026). No new measurements were generated for this study.

### VI.A. Exit is delayed, paced by division and shared between relatives

We found three features that characterise the transition (Fig. 5A–C). First, exit is not memoryless. When each daughter cell is entered into the analysis at its birth (Appendix A), the hazard is negligible for the first *∼* 15 h after 2i withdrawal, and rises to *∼* 0.05 h^*−*1^ by 30–35 h. We compared statistical models of the exit times using a likelihood that accounts for cells that had not exited when last observed (a censored likelihood), ranking the models by the Akaike information criterion (AIC), for which lower values indicate a better fit. This comparison rejects an exponential distribution of exit times, which would correspond to memoryless exit, against every alternative we considered (*Δ*AIC *>* 85), and this conclusion is robust to how commitment events are defined (Fig. S3A). The best Weibull fit has a shape parameter of 2.4, where 1 would indicate memoryless exit, with a 95% interval of 2.0–2.9 from a founder-lineage bootstrap, which resamples whole lineages descended from each founder cell. Second, exit is paced by division. The median commitment occurs 1.5 h after a division (86% within 6 h; Fig. S3B), compared with a median cell-cycle length of 9 h. The commitment hazard falls roughly tenfold from the first two hours after division to mid-cycle, while the division hazard rises from *∼* 6 h and peaks at 14–16 h (Fig. 5B), consistent with the coupling between commitment and division inferred previously (Ruske et al., 2020). The per-cycle probability of committing rather than dividing increases from zero in founder cells to 0.58 by the fifth generation (*χ*^2^ homogeneity *p <* 10^*−*37^), and to 0.50 among cells observed for more than one cycle (Fig. 5A). Third, the latency to commitment is shared between relatives. The exit times of sisters are almost perfectly correlated (*r* = 0.97, 51 pairs), but this largely reflects their shared birth time. When we measured latency from birth instead, the correlation was 0.67 (0.64 for the 40 pairs for which a birth-matched null could be constructed). For comparison, unrelated committed cells matched on imaging date and birth time gave a correlation of 0.10, with a 95% range of *−*0.20 to 0.46 (*p <* 10^*−*3^; Fig. 5C and Fig. S4). First cousins are almost as strongly correlated (0.58, 75 pairs), and the sister of a committing cell commits rather than divides in 83% of cases, compared with 35% for matched unrelated cells. Together, these results suggest that whatever sets the latency persists for at least two generations. This could reflect inheritance or a shared colony environment. With a median of two founder lineages per field of view, these data cannot distinguish the two. Model-independent tests of hereditary structure, which compare the clustering of fates on lineage trees against reshuffling within generations (Allegrezza et al., 2025), could be applied to commit-or-divide fates to test this directly.

**Figure 5.**
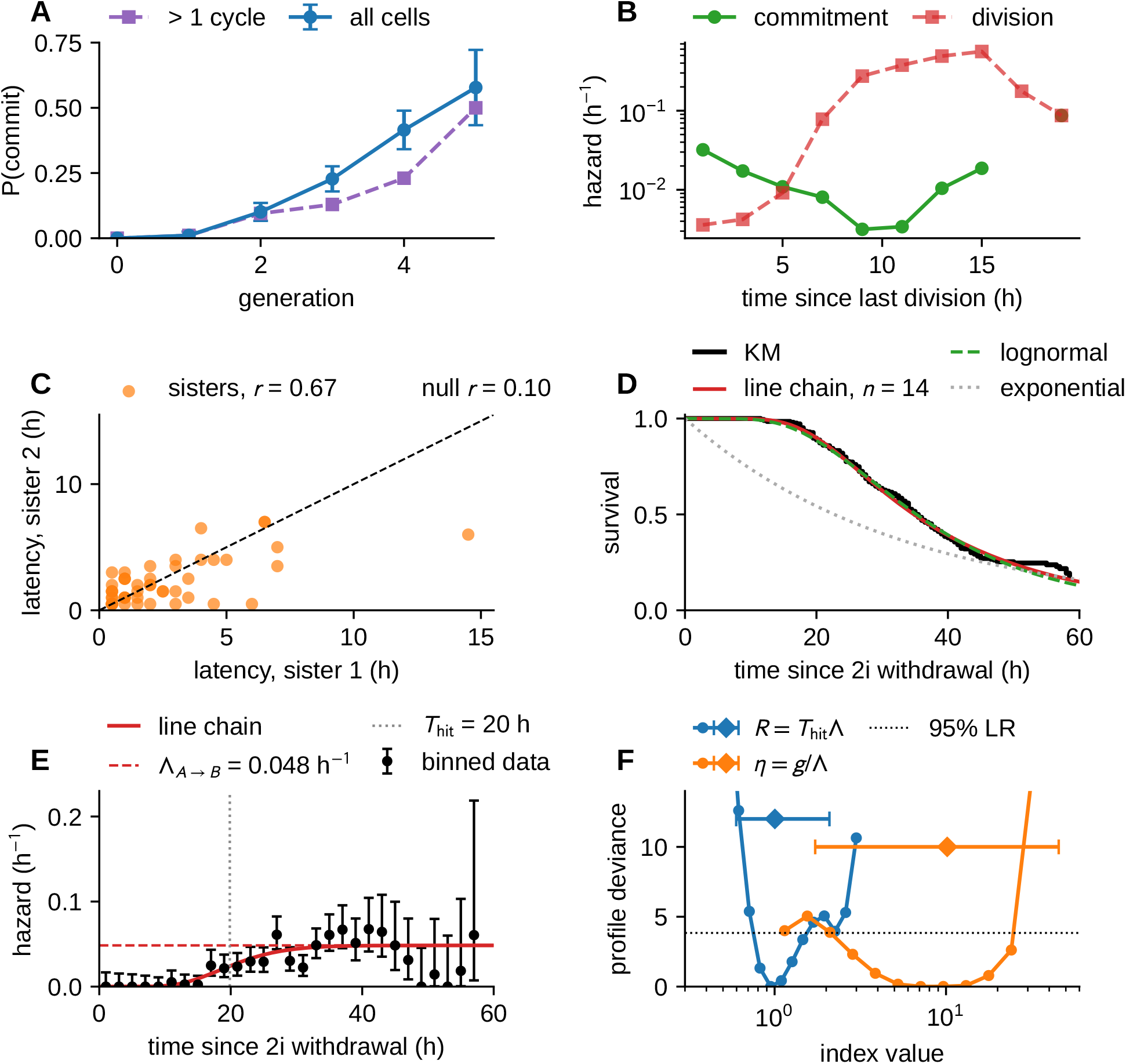
Exit of mouse ESCs from naïve pluripotency. Reanalysis of long-term *Rex1*::GFPd2 imaging (Strawbridge et al., 2026). (A) Per-cycle probability of committing rather than dividing, by generation (all resolved cells, with 95% binomial intervals, and squares for cells observed for more than one cycle). (B) Hazards against time since the last division (log scale), showing that commitment is highest just after division and division peaks late in the cycle. (C) Latency from birth to commitment of sister pairs in which both commit (*r* = 0.67, 51 pairs), against *r* = 0.10 for unrelated committed cells matched on imaging date and birth time. (D) Delayed-entry Kaplan–Meier estimate of survival with the best killed line chain (*n* = 14), the lognormal and the exponential. (E) Hazard estimated as events per cell-hour at risk (occurrence/exposure) in 2-h bins (exact Poisson intervals) with the hazard of the chain, which rises over the lag *T*_hit_ and approaches the quasi-stationary rate Λ_*A→B*_ (dashed). (F) Profile deviance, the loss of fit when an index is held at a given value, minimised over rates and chain length, for *R* = *T*_hit_Λ_*A*_*→*_*B*_ and *η* = *g*/Λ_*A*_*→*_*B*_, with founder-lineage bootstrap medians and 95% intervals (diamonds). The index *R* is identified near 1, whereas *η* is bounded only from above.

A hazard that rises from zero does not, on its own, indicate whether competence acquisition or commitment limits exit, since any killed chain started away from its QSD produces one (Fig. S1A). What matters is how long the rise lasts compared with the persistence time that follows it.

### VI.B. Survival data identify the lag to competence but not the spectral gap

The biological question is whether, after 2i withdrawal, cells spend most of their time becoming able to differentiate, or most of it waiting to do so once able. To estimate these timescales, we fitted the data with the framework’s own killed generators. We used reversible line chains *a*_1_ *⇌ … ⇌ a*_*n*_ *→ B*, with commitment from *a*_*n*_ only and every cell in *a*_1_ at 2i withdrawal, generalising the models of Section II.C (Appendix A). The best-fitting chains fit at least as well as any of the standard statistical distributions we considered (*Δ*AIC = *−*2.7 against the lognormal and *−*36 against the Weibull; Figs. 5D and S3C). Note that the Weibull benchmark here has no delay term, and so differs from the delayed Weibull model of our previous work (Strawbridge and Fletcher, 2026), in which no cells commit before a fitted lag. The fitted chains also agree on their structure, namely a progression through many fast, nearly irreversible steps into the permissive microstate, followed by slow commitment (forward rate 0.69 h^*−*1^, backward rate 0.04 h^*−*1^, and microstate commitment rate *λ* = 0.051 h^*−*1^ for *n* = 14; Fig. 5E). The chain is a phenomenological description of the lag before commitment, and its microstates are not to be identified one-to-one with molecular states.

The quasi-stationary effective commitment rate is Λ_*A→B*_ = 0.048 h^*−*1^ (founder-lineage bootstrap interval 0.036–0.077), corresponding to a mean persistence of *∼* 21 h once cells are competent. The time to first become competent is *T*_hit_ = 20 h (16–29), and the ratio of the two is *R* = 0.96 (0.60–2.09; maximum-likelihood estimates with founder-lineage bootstrap intervals, the bootstrap median of *R* being 1.00). The lag to competence and the subsequent wait are therefore of similar length, with roughly half of the 40-h mean first-passage time spent reaching the permissive set, suggesting that naïve exit is dominated by neither (*R ≈* 1, in the sense defined in Section II.D).

The stability index *η*, by contrast, cannot be estimated reliably from these data (Fig. 5F). Its maximum-likelihood value is *η* = 8.5, with a 95% likelihood-ratio interval (the range of values consistent with the data) of 2.9–24 among chains of up to 20 microstates, and the founder-lineage bootstrap, which allows longer chains, gives 1.7–45. Only the upper end of the range is therefore excluded, reflecting the distinction drawn in Section II.D. In a chain that drifts towards *A*^*\**^, relaxation from the starting state is limited by the time taken to travel along the chain, a time of order *T*_hit_, and not by the spectral gap. Here, 1/*g ≈* 2.4 h, compared with *T*_hit_ *≈* 20 h. Survival data constrain the travel time, through *R*, but not the spectral gap, as expected from the unidentifiability of internal timescales from the survival of an unsorted population (Section IV). Survival curves can therefore tell us how long cells take to become competent, but not how quickly they move among configurations once there, which requires measuring the configurations directly. Two caveats apply to these estimates. First, the chains average over the cell cycle, whereas commitment is coupled to division, so the fitted rates are averages across cell-cycle phases. Second, the estimates assume a single starting microstate and a single permissive microstate.

### VII.C. Knockouts with equal delays leave cells in different internal states

The unidentifiability result of Section IV predicts that perturbations with the same effective commitment rate can leave cells in different internal states, and existing data allow this prediction to be tested at the population level. We used the 73-line Rex1-GFP knockout resource of Lackner et al. (2021), using published differentiation delays based on differentially expressed genes to compare exit timing. The delay accounts for only 26% of the variance in the knockouts’ transcriptional signatures. Moreover, *Rbpj* and *Jarid2*, both delayed by 10.75 h, have weakly anticorrelated residual signatures, meaning the parts of their transcriptional signatures that the delay does not explain (*r* = *−*0.26; Appendix E). However, these bulk data do not resolve the control channel. Commitment-like and competence-like classes of knockouts, defined in advance, do not differ in the size of their residual signatures (exact permutation *p* = 0.07), and they overlap in direction (exact PERMANOVA, a permutation test on the multivariate residuals, *p* = 0.14). A shift between class centroids appears only after excluding knockouts chosen on inspecting the data, and does not survive correction for the choices explored. Together, these results support the premise of unidentifiability but cannot assign the control channel.

### VI.D. Exit from naïve pluripotency provides the time-varying competence that pulsing requires

The pulsing prediction of Section V.B requires competence that varies in time, and there is evidence that this transition provides it. Single-cell transcriptomics has resolved a transient window of heightened susceptibility to lineage-specifying signals as cells exit naïve pluripotency (Semrau et al., 2017), and coupling to division modulates permissive occupancy over the cell cycle (Fig. 5B). In related systems, the duration of a BMP pulse, rather than its concentration, selects fate in pluripotent cells (Camacho-Aguilar et al., 2024), and FGF/ERK signalling is pulsatile both in ESCs (Raina et al., 2022) and in the epiblast of the mouse blastocyst, where its dynamics differ between lineages (Simon et al., 2020). However, none of these studies compares pulsed and sustained signalling at matched dose.

### VI.E. Two experiments would resolve the channel and test the pulsing prediction

Identifying the channel requires measuring the internal state of cells, and not only the timing of their exit. Propositions 4 and 5 suggest a two-tier experimental design. Where a live reporter or surface marker of the permissive set is available, both quantities needed to discriminate the channels can be measured by flow cytometry on rate-matched perturbations. The fraction of cells in the *A*^*\**^ gate gives *π*(*A*^*\**^), and sorting on that gate and timing loss of Rex1-GFP, compared with unsorted cells, gives 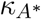. On the solvable network, the commitment cut raises both quantities relative to the competence cut (Fig. 3B,D), and raises permissive occupancy relative to a rate-matched mixing cut, itself additionally separated by the gap of *Q*_*A*_ (Fig. S2D). Where no such marker exists, *A*^*\**^ must itself be inferred. In this case, we propose profiling a 2i-withdrawal time course (for example, at 0, 12, 24, 36 and 48 h after 2i withdrawal) by single-cell RNA-seq with metabolic labelling (Qiu et al., 2020), in the RC9 (*Rex1*::GFPd2; Cas9) line used for the knockout resource (Lackner et al., 2021). The experiment would carry a rate-matched pair of guides, with two guides per gene and *∼*100 cells per guide. We would pair *Tcf7l1*, a naïve-network node whose loss reinforces naïve circuitry and is hypothesised here to restrict access to the permissive set through *Q*_*A*_, with *Rbpj*, the Notch effector whose loss lowers the commitment gate (a commitment cut) (Lackner et al., 2021). Both knockouts delay exit and can be titrated to a common Rex1-GFP rate. Metabolic labelling distinguishes new from old RNA, providing the directionality needed to reconstruct *Q*_*A*_. The macrostates, *A*^*\**^ and *π*(*A*^*\**^) then follow by absorbing Markov-chain analysis (Lange et al., 2022; Weinreb et al., 2018). We predict that, at matched rate, cells will accumulate in the permissive set behind the lowered gate of the commitment cut (high *π*(*A*^*\**^)), but not under the mixing cut (low *π*(*A*^*\**^)), and the mixing cut should in addition alter the reconstructed *Q*_*A*_. If rate-matched cuts of the two classes instead share *π*(*A*^*\**^) and sorted first-passage times, the three-way decomposition is not realised in this system. To our knowledge, no time-resolved single-cell CRISPR screen of this transition exists. The closest, in human induced pluripotent stem cells (Ishikawa et al., 2023), perturbs pluripotency-network factors but lacks a signalling member, and so does not provide the matched pair required.

The pulsing prediction can be tested with the same reporter. We predict that commitment pulses timed to measured peaks in competence will commit more cells than sustained signalling of equal integrated dose, and that the advantage will rise from zero as the depth of competence modulation increases. Because the size of the advantage depends on that depth (Section V.B), the most informative design varies the depth of competence modulation, for example by partially synchronising the cell cycle, instead of comparing a single pair of protocols. The absence of an advantage where competence demonstrably varies, or an equal advantage where it does not, would falsify this prediction.

## VII. Discussion

In this study, we developed a framework that describes the stability of a cell identity by its effective commitment rate, and used it to ask how that stability can be controlled. We found that the sensitivity of the effective commitment rate separates contributions from mixing, competence, and commitment. At a fixed competence profile, the elasticities to uniform scaling of commitment rates and microstate transition rates sum to one. These elasticities locate the model on a scale from commitment-limited to accessibility-limited. Interventions with equal effective commitment rates cannot be distinguished from the survival of an unsorted population, but can be distinguished by the occupancy of the permissive set or by first passage from sorted cells. When the channels are treated as inputs, pulsed commitment signalling helps, in the models we studied, only when competence varies in time. Since many developmental signals, ERK among them, are delivered in pulses (Simon et al., 2020; Raina et al., 2022; Mulas et al., 2024), this suggests that pulsing may serve to match signal to fluctuating competence, in addition to arising from feedback within signalling pathways, a hypothesis that the experiments of Section VI.E could test. For the exit of mouse ESCs from naïve pluripotency, the fitted chains give comparable estimates for the mean competence lag and the subsequent mean wait to commitment.

### VII.A. Development, homeostasis and regeneration may be operating points of one system

Our framework summarises the stability of an identity by a small number of quantities, namely the effective commitment rate, the shares of control held by commitment and by movement among microstates, and the timescale indices *η* and *R*. Development, homeostasis and regeneration can then be viewed as different operating points of the same system, instead of as distinct mechanisms. Development maintains high competence and promotes commitment, so that identities are intentionally transient. For naïve exit, an in vitro transition, we estimated a lag to competence comparable to the persistence that follows it (*R ≈* 1), with commitment coupled to division. Homeostasis would correspond to low competence and mixing much faster than effective commitment (*η »* 1), with identity stabilised by restricting access to permissive configurations, so that transient fluctuations in signalling rarely drive a change of fate. Regeneration and repair would correspond to time-dependent modulation of the same parameters. Injury-induced signals that transiently enlarge or expose the permissive set, or increase microstate commitment rates, could lower stability for long enough for cells to contribute to repair, after which the baseline parameters are restored. This view is consistent with the existence of competence windows during regeneration.

### VII.B. A stable identity need not sit in a deep basin

In a landscape picture, stability is associated with the depth of a basin. In our framework, stability is measured by the effective commitment rate, set by the accessibility of permissive microstates and by the kinetics of commitment once they are reached. An identity can therefore be highly stable without a deep basin, provided that permissive microstates are rarely occupied or poorly connected. Even in deterministic Boolean models, attractor states can be less robust to perturbation than the basins that lead to them (Bavisetty et al., 2025), so the size of a basin is a poor guide to the persistence of the identity it contains. Landscapes for gene regulatory networks, which operate far from equilibrium, are in any case metaphors, since no true potential exists in general (Moris et al., 2016). Where the microstate dynamics are approximately reversible, the QSD takes a Boltzmann-like form, with occupancy falling exponentially with an effective energy, and an effective equilibrium landscape re-emerges. Where it does not, landscape-and-flux theories retain a landscape by adding a circulating flux to the quasi-potential (Wang, 2015), and recent work has derived such landscapes statistically from data (Sáez et al., 2022; Raju and Siggia, 2023). Our framework complements these approaches by asking which inputs move a cell between identities, and on what schedule. The separation of channels also helps to explain observations that are difficult to interpret with a static landscape, such as strong signals that fail when competence is restricted, and predicts when pulsed exposures should outperform sustained exposures of equal dose (Sections V.B and VI.E).

### VII.C. Limitations and outlook

Our framework has several limitations. First, it assumes that cell states can be coarse-grained into microstates whose dynamics are Markovian, so that the current microstate carries all the information needed to predict subsequent transitions. Estimating *Q*_*A*_, *A*^*\**^ and *λ* from data is the natural next step, and existing single-cell pipelines provide a route. A nearest-neighbour graph of cells, made directional by metabolic labelling or RNA velocity, approximates *Q*_*A*_, spectral coarse-graining, which groups cells by the slowest modes of the dynamics, identifies macrostates and the permissive set, and absorption probabilities give commitment rates (Lange et al., 2022; Weinreb et al., 2018). Propositions 4 and 5 show that *λ* need not be estimated separately once *Q*_*A*_ and either the occupancy or sorted first-passage times are available. Proposition 1 provides the gradient needed for design problems at the cost of computing one principal eigenpair, independently of the number of parameters. The models studied here are deliberately minimal, whereas graphs inferred from data would comprise thousands of microstates.

Second, all our test systems are reversible, whereas biological microstate graphs, driven by energy-consuming chromatin remodelling and signalling, are generally not. For non-reversible generators, the hazard can overshoot Λ_*A→B*_ during the first few relaxation times, by up to 36% in our examples, and the error in approximating *T*_*A→B*_ by 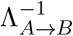 roughly doubles at matched *η* (Appendix D). Since velocity-based reconstructions are directional by design, this is likely to be the typical situation in data. The sensitivity decomposition, the elasticity bounds and the identifiability results require only irreducibility, and so do not depend on detailed balance. The identification of relaxation with the inverse spectral gap, and the convexity results for mixing control, do depend on it. Conclusions about which channel to target therefore hold whether or not the underlying dynamics are driven by energy-consuming processes, whereas statements about how quickly cells forget their starting configuration should be treated with more caution. Finally, our models treat cells as independent, whereas in the data exit is coupled to division and correlated between relatives. Coupling the killed generator to a branching process is a natural extension, and would allow the inherited component of latency to be estimated directly.

## ACKNOWLEDGMENTS

S.E.S. was supported by a Sir Henry Wellcome Postdoctoral Fellowship (224070/Z/21/Z) and a University of Sheffield Strategic Research Fellowship in the Physics of Life and Quantitative Biology.

A.G.F. was supported by the Engineering and Physical Sciences Research Council (EP/W024144/1). The funders had no role in study design, analysis, decision to publish, or preparation of the manuscript. For the purpose of open access, the authors have applied a Creative Commons Attribution (CC BY) licence to any Author Accepted Manuscript version arising from this submission. Generative AI (Anthropic’s Claude) was used to assist with editing the manuscript text and associated code. All AI-assisted text and code were reviewed, verified and edited by the authors, who take full responsibility for the content and conclusions of the work.

## AUTHOR CONTRIBUTIONS

S.E.S. and A.G.F. contributed equally to Conceptualization, Formal analysis, Funding acquisition, Methodology, Software, Validation, Visualization, Writing – original draft, Writing – review & editing.

## DATA AND CODE AVAILABILITY

This study generated no new experimental data. Code to reproduce every numerical result and figure is openly available at https://github.com/stanleystrawbridge/strawbridge_and_fletcher_2026_mixing. The script `cell_state_numerics.py` generates Figs. 1–4, S1, S2 and S5, and verifies Proposition 1, Corollary 2 and Propositions 4–5 numerically. The script `cell_state_figures.py` performs the single-cell and bulk reanalyses (Figs. S4 and S6), `cell_state_stability_fit.py` fits the killed chains (Fig. S3), and `fig_naive_exit.py` generates Fig. 5. The single-cell imaging data are from the repository accompanying Strawbridge et al. (2026) (commit `8bdd129`), the knock-out transcriptomes are from GEO (GSE145653) and the differentiation delays from Dataset EV2 of Lackner et al. (2021).

## A. Methods

### Numerical methods

Principal eigenpairs of killed generators were computed by dense eigen-decomposition, survival curves by spectral decomposition of *M* ^*⊤*^, and mean first-passage times by solving *−MT* = **1**. Interventions were matched in effective commitment rate by bisection to a tolerance of 10^*−*14^. The optimal commitment input was computed from the Pontryagin conditions by a damped forward–backward sweep (60 iterations, damping 0.35, fourth-order Runge–Kutta on 4001 time points), the bang–bang input at each iteration saturating where the switching function is largest subject to the dose constraint. The quasi-static argument of Section V.B shows that a constant input is optimal under stationary competence only among slowly varying inputs. A general result would require second-order conditions for so-called singular arcs, stretches of time on which the optimal input is neither fully on nor fully off, and we therefore establish it only for the models studied. Schrödinger bridges were computed by Sinkhorn iteration on the reference kernel exp(*T Q*_*A*_) to a tolerance of 10^*−*14^, with endpoint distributions softened to 0.05 for numerical conditioning. Every numerical value quoted in Sections II–V is printed by cell_state_numerics.py, which also verifies Proposition 1, Corollary 2, the Q-process identity and Propositions 4–5 numerically.

### Survival analysis of single-cell exit

Tracked-lineage records were taken from the Strawbridge & Corridori repository (commit 8bdd129, N2B27 wells only). A cell was scored as committed at the first frame after its last above-threshold frame, provided its final frame lay below threshold, the threshold being the upper one-sided 95% *t*-bound of log_10_ fluorescence in the same-day 2i negative control. Time is measured from 2i withdrawal. Because daughter cells exist only from their birth, each cell enters the risk set at its first frame and leaves at commitment (event) or at division, death, loss or the end of imaging (right-censored). Survival was estimated by the Kaplan–Meier estimator with delayed entry and the hazard by occurrence/exposure rates in 2-h bins with exact Poisson intervals. Parametric models were fitted by maximising the left-truncated, right-censored likelihood Σ_events_ log *f*(*t*_*i*_)+ Σ_censored_ log *S*(*t*_*i*_)*− Σ* _all_ log *S*(*e*_*i*_), with *e*_*i*_ the entry time, and compared by AIC. Uncertainty was assessed by a cluster bootstrap over founder lineages (107 lineages, 200 replicates), which respects the dependence between relatives. Each field of view is identified by imaging date and file name, since file names recur across the two imaging dates.

### Killed-chain fits and the stability index

First-passage data were fitted with reversible line chains on *n* microstates, with forward rate *a*, backward rate *b* and commitment rate *λ* from the terminal microstate only, every cell starting in the first microstate at 2i withdrawal. The homogeneous chain has three parameters for any *n*, and *n* = 2, …, 20 was selected by AIC (chains with free rates on every edge, *n* = 2–4, fitted worse). Because frames are 30 min apart, commitment was treated as interval-censored between the last frame above and the first frame below threshold, together with right censoring and delayed entry, and the lognormal and Weibull benchmarks were refitted with the same likelihood. Survival on the frame grid was computed by propagating the initial distribution with exp(*Δt M*). From each fit we report Λ_*A→B*_ and the gap *g* of *Q*_*A*_, the mean time *T*_hit_ for the closed chain to first reach the permissive microstate, *R* = *T*_hit_Λ_*A→B*_, and *η* = *g*/Λ_*A→B*_. Profile likelihoods for *R* and *η* were minimised over the rates and over every *n* within 10 AIC units of the best. Because *n* is weakly identified at large *n*, confidence intervals were also obtained by a founder-lineage bootstrap (200 replicates) in which *n* was re-selected in every replicate, which additionally accounts for the correlation between relatives. Line chains cannot produce *η <* 1, so the lower limit on *η* is set by the model family rather than the data.

### Correlation between relatives

For sister pairs (sharing a parent) and first-cousin pairs (sharing a grandparent but not a parent) in which both cells commit, we correlated the latency from birth to commitment. Absolute commitment times are not used, because relatives born together share their time since 2i withdrawal and the commitment hazard depends on that time. The null distribution was built by replacing each sister pair with two committed cells from different founder lineages, imaged on the same date and born within 0.5 h of the pair, and recomputing the correlation (5000 draws, using the 40 of 51 pairs with at least two such cells and computing the observed statistic on the same pairs). This null retains the shared clock and the selection on both cells committing, so any excess correlation is attributable to relatedness or to a shared local environment. Fate concordance compares the probability that the sister of a committing cell commits rather than divides with the corresponding probability for birth-matched unrelated cells.

## B. Standard derivations

These results are standard for continuous-time Markov chains with absorption and are included for completeness.

### B.1. Exit hazard

We retain the notation of the main text, with *X*_*t*_ the microstate at time *t* and *τ*_*A→B*_ the first-passage time to *B*. The exit hazard is the instantaneous rate of commitment conditioned on survival within *A*,

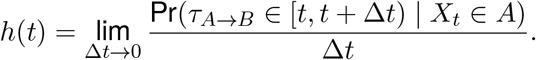

Writing *p*_*i*_(*t*) = Pr(*X*_*t*_ = *a*_*i*_, *τ*_*A→B*_ *> t*), the survival-conditioned occupancy of *a*_*i*_ is 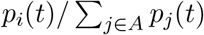, and commitment from *a*_*i*_ within *Δt* has probability *λ*_*i*_*Δt* + *o*(*Δt*). Averaging over microstates and letting *Δt →* 0,

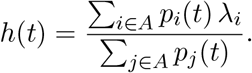

Because *λ*_*i*_ = 0 for *a*_*i*_ *∈*/ *A*^*\**^, only permissive microstates contribute.

### B.2. Quasi-stationary distribution and effective commitment rate

Conditioning the killed process on non-absorption defines the quasi-stationary distribution

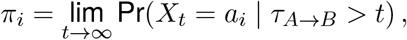

when the limit exists (Darroch and Seneta, 1967). The killed dynamics 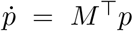, with substochastic generator *M* = *Q*_*A*_ *−* diag(*λ*), have a principal (least-negative) eigenvalue *−*Λ_*A→B*_ with corresponding left eigenvector *π*, normalised as a probability distribution,

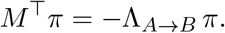

Under a separation of timescales between mixing within the identity and commitment, transient contributions from the subdominant spectrum decay and the survival probability decays exponentially, 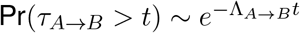, and escape is then asymptotically a Poisson process (Freidlin and Wentzell, 2012; Bovier and den Hollander, 2015) with rate

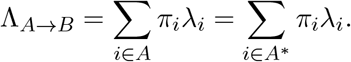

Equivalently, evaluating the main-text hazard at the quasi-stationary distribution gives the time-independent rate *h* = Σ_*i*_ *π*_*i*_*λ*_*i*_ = Λ_*A→B*_. The mean first-passage time (MFPT) from a quasistationary initial condition is the inverse of this rate, 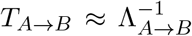, with relative corrections of order Λ_*A→B*_/*g*, where *g* is the spectral gap of *Q*_*A*_. These corrections are negligible precisely in the metastable regime Λ_*A→B*_ *« g*.

### B.3. Mean first-passage times in the two-state model

Let *T*_*N*_ and *T*_*P*_ be the mean times to absorption from *N* and *P*. From *N* the only transition is to *P* at rate *α*, so *T*_*N*_ = 1/*α* + *T*_*P*_ . From *P* the next event occurs after a mean time 1/(*β* + *λ*) and is a return to *N* with probability *β*/(*β* + *λ*), so *T*_*P*_ = 1/(*β* + *λ*) + *βT*_*N*_ /(*β* + *λ*). Hence

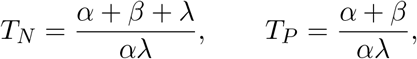

and averaging over the closed-chain stationary distribution *π*^0^ = (*β, α* /(*α* + *β*) gives

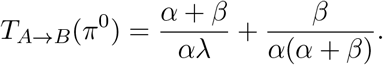

The first term is the inverse of the leading-order effective commitment rate *αλ*/(*α* + *β*), and the second an *O*(1) correction that is negligible when *λ « α* + *β*.

## C. Proofs

The results below are not standard. All require *Q*_*A*_ irreducible on *A* and *A*^*\**^ *≠ ∅*, and none requires reversibility.

### C.1. Proof of the sensitivity decomposition

Let *M* = *Q*_*A*_ *−* diag(*λ*) be irreducible with principal eigenvalue *θ* = *−*Λ_*A→B*_, left eigenvector *π* and right eigenvector *ϕ*, normalised by Σ_*i*_ *π*_*i*_ = 1 and *π*^*⊤*^*ϕ* = 1. Irreducibility makes *θ* simple, so it depends analytically on the entries of *M* in a neighbourhood.

Differentiating *Mϕ* = *θϕ* and contracting on the left with *π*^*⊤*^,

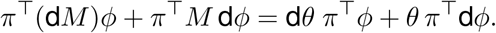

Since *π*^*⊤*^*M* = *θπ*^*⊤*^, the terms in d*ϕ* cancel, and with *π*^*⊤*^*ϕ* = 1,

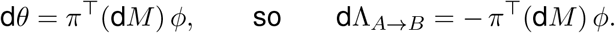

This is the non-symmetric form of the Hellmann–Feynman relation, in which the eigenvector perturbation drops out because the eigenvalue is stationary in it.

Now d*M* = d*Q*_*A*_ *−* diag(d*λ*), so

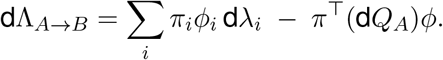

For the second term, d*Q*_*A*_ must preserve zero row sums, so d*q*_*ii*_ = *−* Σ_*j≠i*_ d*q*_*ij*_ and

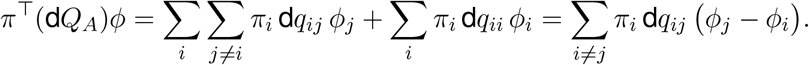

Substituting gives the unconstrained partial derivatives stated in Proposition 1. Writing *λ* = *Uℓ* gives d*λ*_*i*_ = *ℓ*_*i*_ d*U* + *U* d*ℓ*_*i*_ with Σ_*i*_ d*ℓ*_*i*_ = 0, and splitting the first sum accordingly yields the three-term form.

The probabilistic reading of *ϕ* follows from the spectral expansion of the killed semigroup, 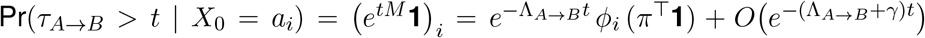, where *γ >* 0 is the gap between *−*Λ_*A→B*_ and the real part of the subdominant eigenvalue of *M*. Since *π*^*⊤*^**1** = 1, *ϕ*_*i*_ is the asymptotic survival amplitude of a cell started in *a*_*i*_.

### C.2. Proof of identifiability from the resolved quasi-stationary distribution

The quasi-stationary distribution satisfies *M* ^*⊤*^*π* = *−*Λ_*A→B*_*π* with 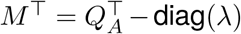. Componentwise,

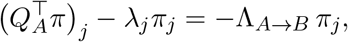

and since *π*_*j*_ *>* 0 for all *j* by Perron–Frobenius,

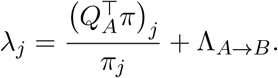

The map Λ_*A→B*_ *1↦ λ* is thus an affine one-parameter family of uniform shifts, and no further information is available from the eigenvector equation alone, since summing *π*_*j*_*λ*_*j*_ over *j* and using *Q*_*A*_**1** = 0 returns Σ_*j*_*π*_*j*_ *λ*_*j*_ = Λ_*A→B*_, which is the defining identity rather than an additional constraint. If 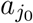 is known to be non-permissive then 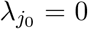 fixes 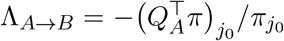 and hence *λ* throughout. Any second non-permissive microstate must return the same value, giving |*A\A*^*\**^|*−*1 over-identifying restrictions.

### C.3. Proof of identifiability from first passage

Since *Q*_*A*_ is irreducible and *A*^*\**^ *≠ ∅, M* is invertible and every *T*_*i*_ is finite and positive. The mean first-passage times satisfy the backward equation *−MT* = **1**, that is *λ*_*i*_*T*_*i*_ *− Q*_*A*_*T* _*i*_ = 1 for each *i*, which rearranges to the stated expression. Each *λ*_*i*_ is thus an explicit function of (*Q*_*A*_, *T* ), so the map *λ 1 ↦ T* at fixed *Q*_*A*_ is injective, and distinct *λ* give distinct *T* . For non-permissive microstates the numerator vanishes identically, 1 + (*Q*_*A*_*T* )_*i*_ = 0, which supplies |*A \ A*^*\**^| consistency checks when *A*^*\**^ is known.

### C.4. Proof of the elasticity identity and bounds

Scaling every rate by *c >* 0 scales *M*, and hence its principal eigenvalue, by *c*, so Λ_*A→B*_ is homogeneous of degree one in (*q*_*ij*_, *λ*_*i*_) jointly. Euler’s theorem gives 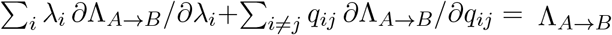 With *λ* = *Uℓ* at fixed 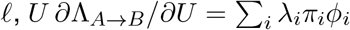 by Proposition 1, and dividing by Λ_*A→B*_ gives *ε*_*U*_ + *ε*_*Q*_ = 1. For the bounds, the dominant eigenvalue of an essentially non-negative matrix is a convex function of its diagonal (Cohen, 1981). Hence Λ_*A→B*_(*U*) = *−λ*_max_ *Q*_*A*_ *− U* diag(*ℓ*) is concave in *U*, with Λ_*A→B*_(0) = 0 because *Q*_*A*_ has zero row sums. Concavity with Λ_*A→B*_(0) = 0 gives Λ_*A→B*_(*U*) *≥ U ∂*Λ_*A→B*_/*∂U*, so *ε*_*U*_ *≤* 1, and *∂*Λ_*A→B*_/*∂U* = Σ _*i*_ *ℓ*_*i*_*π*_*i*_*ϕ*_*i*_ *>* 0 gives *ε*_*U*_ *>* 0. The bounds on *ε*_*Q*_ follow from the identity. That (*π*_*i*_*ϕ*_*i*_) is the stationary distribution of the Q-process follows by direct substitution, since the transformed generator 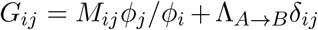 has zero row sums because 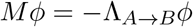, and 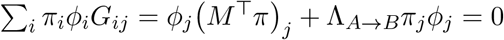.

### C.5. Proof of the size of a stability class

By Proposition 1, *∂*Λ_*A→B*_/*∂λ*_*i*_ = *π*_*i*_*ϕ*_*i*_ *>* 0 for every *i*, since both eigenvectors are strictly positive, so *∇*Λ_*A→B*_ *≠* 0 everywhere on Θ. The implicit function theorem then makes every level set of Λ_*A→B*_ a smooth embedded hypersurface of codimension one, with normal *∇*Λ_*A→B*_. The asymptotic survival curve, Λ_*A→B*_ and the mean first-passage time from quasi-stationarity, 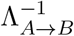,are functions of Λ_*A→B*_ alone, and so are constant on each class.

### C.6. Monotonicity of the effective commitment rate in symmetric edge weights

For an edge of weight *w*_*ij*_ = *q*_*ij*_ = *q*_*ji*_, the two mixing terms of Proposition 1 combine to

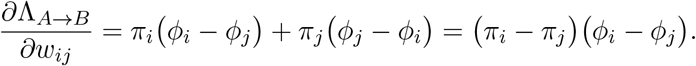

When *M* is symmetric its left and right principal eigenvectors coincide up to normalisation, so *π ∝ ϕ* and the product is non-negative. Hence Λ_*A→B*_ is non-decreasing in every edge weight as well as in every microstate commitment rate.

## D. Departures from the reversible picture

The main text uses reversible generators throughout. Biological microstate dynamics, however, are driven by energy-consuming processes such as chromatin remodelling and signalling, and so need not be reversible. To quantify what is lost when that assumption fails, we compare the eleven-microstate reversible network of Fig. 1B with two non-reversible generators on the same number of microstates, namely a uniform directed cycle (*i → i* + 1 at rate 1, *i → i −* 1 at rate 0.02) and a heterogeneous directed cycle with forward rates drawn uniformly from [0.3, 2]. In each case the three microstates furthest along the cycle constitute *A*^*\**^, with *λ* = 0.15, and the process is started from a point mass far from the permissive set. The three systems have comparable stability index (*η* = 4.9, 4.2 and 4.2 respectively), so they are compared at matched timescale separation.

The two non-reversible mechanisms separate cleanly (Fig. S5). The uniform directed cycle is a circulant matrix and therefore *normal*, in that its departure from normality, measured by the Henrici index 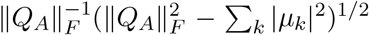 with *µ*_*k*_ the eigenvalues, is zero to machine precision. It nonetheless has strongly complex eigenvalues (max_*k*_ | Im *µ*_*k*_| = 0.97), and the exit hazard overshoots the quasi-stationary rate by 36% before ringing down to it. The heterogeneous cycle is genuinely non-normal (Henrici index 0.48) and shows transient amplification of ‖ *e*^*tM*^ *‖*_2_ by a factor of 1.13 above the asymptotic decay, with a hazard overshoot of 27%.

Two conclusions follow. First, the qualitative signature of the quasi-stationary reduction in Fig. S1A – a hazard rising monotonically from zero to Λ_*A→B*_ – is a property of reversibility, not of metastability. Under either non-reversible mechanism the hazard is non-monotone and the survival curve departs visibly from a single exponential over the first mixing times. Second, the quantitative cost is material but bounded. The relative error in 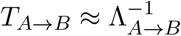 rises from 6.3% for the reversible network to 14.2 % and 16.7% for the two cycles at matched *η*, so the reduction degrades by roughly a factor of two rather than failing outright. The asymptotic content of the framework is unaffected – *π, ϕ*, Λ_*A→B*_ and Propositions 1–5 require only irreducibility – but the identification of *τ*_mix_ with 1/*g*, and any inference that reads a lag time as a mixing time, should be treated with caution when the reconstructed generator is strongly non-reversible. Since RNA-velocity and metabolic-labelling reconstructions are directional by design, this is the expected situation in data rather than an edge case.

## E. The bulk knockout resource

### Data and commitment-rate proxy

The saturating Rex1-GFP knockout resource of Lackner et al. (2021) (see also Leeb et al., 2014) comprises 73 isogenic knockout lines differentiated in parallel, each with a transcriptome in 2i and at N24 (GEO GSE145653), together with a 2-hresolved wild-type differentiation time course. Projecting each knockout’s N24 transcriptome onto that time course gives a differentiation delay *Δt*_KO_ in hours (Dataset EV2 of the original study), which we take as a proxy for the effective commitment rate. Under a proportional-hazards reading, Λ_KO_/Λ_WT_ *≈* (24 *− Δt*_KO_)/24. The resource reports two delays, one from differentially expressed genes and one from naïve-marker expression. We use the former and repeat every analysis with the latter. Two limitations bound what this test can establish. There is no per-knockout survival curve, so Λ_KO_ is read from a delay rather than an asymptotic slope. Moreover, a delay conflates a longer lag with a lower commitment rate, the very distinction the framework draws, so the bulk test reaches only the population-mean form of the prediction.

### Classification

The channel classification was fixed before any residual was examined. From the resource’s signalling footprints and known biology, the commitment-like (signalling) class is *{Rbpj, Fgfr1, Fgfr2, Ptpn11, Raf1, Mapk1, Etv5, Tsc2}* and the competence-like (chromatin and naïve-network) class is *{Eed, Suz12, Jarid2* (PRC2), *Mbd3, Mta3* (NuRD), *Tet1, Kdm6a, Tcf7l1, Zfp281, Trim71}. Csnk1a1* is excluded for a passage-adaptive proliferation confound, commitment being coupled to division in this system (Ruske et al., 2020). *Tcf7l1* defines the Wnt footprint yet is mechanistically a naïve-network node; we place it with competence but note the ambiguity. The class labels reflect a biological grouping fixed before analysis. In the coordinates of Section III.B, several competence-like knockouts, *Tcf7l1* among them, are expected to act on the accessibility of the permissive set, and hence on *Q*_*A*_, and not on *ℓ* alone.

### Analysis

Counts were normalised to log_2_ counts per million, and each knockout’s N24 signature was taken as its mean log fold change against batch-matched RC9 wild-type controls over the 2000 most variable expressed genes. Signatures of all 73 knockouts were regressed on a quadratic in the delay, splitting each into an on-trajectory component and an off-trajectory residual, and the residual norm was then adjusted linearly for delay. Residual magnitude was compared between classes with a one-sided exact permutation test over all 43,758 relabellings of the 18 classified knockouts, and residual direction was compared with an exact PERMANOVA (Euclidean distance, pseudo-*F*) over the same relabellings, together with a test of within-class dispersion to separate a shift in location from a difference in spread.

### Results

The delay accounts for 26% of the variance in knockout signatures (20% with the naïve-marker delay), and for the median knockout, 8% of its signature lies on the delay trajectory. *Rbpj* (commitment-like) and *Jarid2* (competence-like) share a delay of 10.75 h and have weakly anti-correlated residuals (*r* = *−*0.26); with the naïve-marker delay the two are no longer matched (19.0 against 13.25 h). Equal delay does not fix internal state. The competence-like residual is larger than the commitment-like residual on average, but not significantly so (exact *p* = 0.072, or 0.070 with the naïve-marker delay), and the difference is carried by the PRC2 lines, which the resource flags for disproportionate global deregulation, and without them *p* = 0.33 (0.33). In residual space the classes overlap (PERMANOVA *p* = 0.14, or 0.09 with the naïve-marker delay), with centroids separated by less than the typical distance of a knockout from its class centroid (ratio 0.65). Excluding PRC2 yields a centroid shift (PERMANOVA *p* = 0.03, or 0.02 with the naïve-marker delay) that is not a dispersion artefact, the two classes being equally spread once PRC2 is removed. Excluding *Rbpj* and *Tcf7l1*, the pair proposed for the prospective experiment, makes the magnitude difference nominally significant (*p* = 0.04), again carried by PRC2 (*p* = 0.22 once PRC2 is also excluded), and leaves the directional test null while PRC2 is included (*p* = 0.21). With both PRC2 and the pair excluded the centroid shift strengthens (PERMANOVA *p* = 0.009, 6 against 7 knockouts). Every exclusion here was chosen after inspecting the data, and across the 8 combinations of delay definition and class set reported in `bulk_summary.csv` the smallest PERMANOVA *p* becomes 0.07 after Bonferroni correction over those combinations alone. The directional separation among knockouts with clean mechanisms is therefore a hint worth testing prospectively, not a finding. For genetic knockouts, moreover, distinct perturbations matched in rate trivially disturb distinct genes, so the existence of a residual is uninformative. What the framework needs is a residual that resolves into competence and commitment channels, and the bulk data do not deliver this. These analyses are implemented in `cell_state_figures.py`.

## F. Designing stability and minimum-effort steering

This appendix extends the control formulation of Section V.A to two design questions, namely which admissible rates best maintain or destabilise an identity, and how a population can be steered to a target distribution with least disruption.

### F.1. Optimal protocols push independently tunable rates to their limits

Maintaining an identity against noise amounts to choosing admissible rates that minimise Λ_*A→B*_ = *−λ*_max_(*M*), while inducing differentiation amounts to maximising it. We asked whether these two problems are equally hard. The dominant eigenvalue of an essentially non-negative matrix is convex in its diagonal entries (Cohen, 1981), so Λ_*A→B*_ is concave in the microstate commitment rates for any generator. For symmetric transition rates, *M* is symmetric and the same holds for the edge weights, as in the design of fastest-mixing Markov chains (Boyd et al., 2004). Destabilisation over a convex set of admissible rates is then a convex optimisation problem, whereas stabilisation minimises a concave function, whose optimum lies at a vertex of a polytope of admissible rates. In practice, this asymmetry is weaker than it might appear. Because Λ_*A→B*_ increases with every microstate commitment rate (Proposition 1) and, for symmetric rates, with every edge weight (Appendix C.6), both optima lie at corners when each rate is bounded independently. The two problems differ in difficulty only under coupled constraints, such as a shared budget, and we are not aware of a general result for non-symmetric parameterisations of the graph. Put simply, when each rate can be tuned independently, the best protocol for either goal pushes every rate to one of its limits, and finding that protocol becomes hard only when rates compete for a shared resource. We note that this concerns the design of interventions, not the biology, since a cell maintains its identity by operating at a particular set of parameters, not by solving an optimisation problem.

### F.2. Minimum-effort steering reweights transitions that cells already make

Directed interventions, such as differentiation protocols, reprogramming cocktails and regenerative cues, can be viewed as steering the microstate distribution from an initial state *ρ*_0_ to a target *ρ*_*T*_ at least cost. Scheduling interventions on a Markov model to steer a population towards a target has precedent in the design of sequential antibiotic therapies, where the order of drugs is chosen to steer evolution on a fitness landscape away from resistance (Nichol et al., 2015). If cost is measured by the relative entropy of the controlled process with respect to the unperturbed process, a measure of how far the controlled dynamics depart from the natural ones, this is the Schrödingerbridge problem (Chen et al., 2021). The same cost renders optimal control of stochastic reaction networks linearly solvable (Horiguchi and Kobayashi, 2025). On a finite graph, its solution is a Doob transform of the reference generator that reweights the existing transition rates by time-dependent factors. These factors are obtained by a forward–backward (Sinkhorn) iteration, so the optimal intervention amounts to mixing control with an explicit, minimum-effort schedule (Fig. S2A,B). For a differentiation protocol, the bridge therefore gives the least disruptive way of adjusting the transitions that cells already make, so that a chosen distribution of cells is reached by a chosen time. Because relative entropy is infinite on paths that the unperturbed process never takes, no finite-cost control can create a transition absent from *Q*_*A*_. This follows from the choice of cost, and defines the scope of the approach. The cost is suited to interventions that modulate transitions cells already make, for example by altering signalling, mechanics or metabolic state, and it penalises forcing cells along trajectories they rarely explore, since such forcing in practice brings metabolic burden, toxicity or off-target epigenetic change. It excludes interventions whose purpose is to open new routes, such as forced expression of a transcription factor that a cell does not otherwise express. Costs that make creating a transition expensive, instead of impossible, would extend the framework to direct reprogramming. The Q-process of Section III.C is also a Doob transform of the killed generator, by *ϕ*, so long-term survival and minimum-effort steering correspond to the same construction, conditioning the process either on never committing or on arriving at *ρ*_*T*_ .

## SUPPLEMENTAL FIGURES

**Figure S1.**
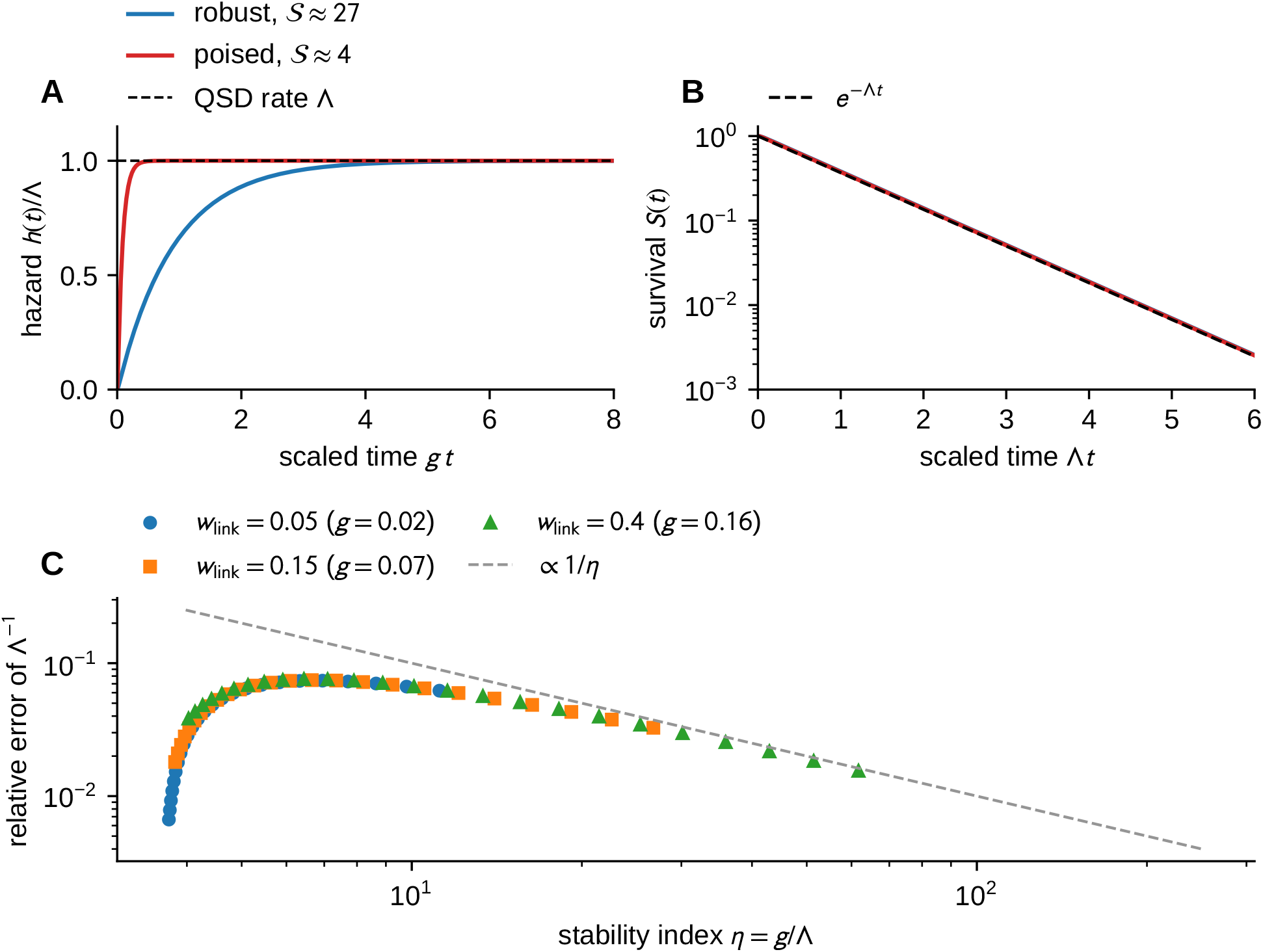
Validation of the quasi-stationary reduction. Eleven-state network of Fig. 1B, commitment from the pocket, started from the bulk microstate farthest from the bridge. (A) The normalised hazard rises from zero and relaxes to Λ_*A→B*_ within the relaxation time 1/*g*, faster for a robust (*η ≈* 27, *λ* = 0.01) than for a poised (*η ≈* 4, *λ* = 0.8) identity. (B) Survival collapses onto 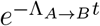 once initial conditions wash out. (C) The relative error of 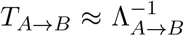 falls as 1/*η* for *η ≳* 10 across network geometries (*w*_link_ *∈ {*0.05, 0.15, 0.4*}*) and effective commitment rates spanning two decades. It peaks near *η≈*6, and the low-*η* branch, where a single slow crossing sets both quantities, is not evidence for the reduction.

**Figure S2.**
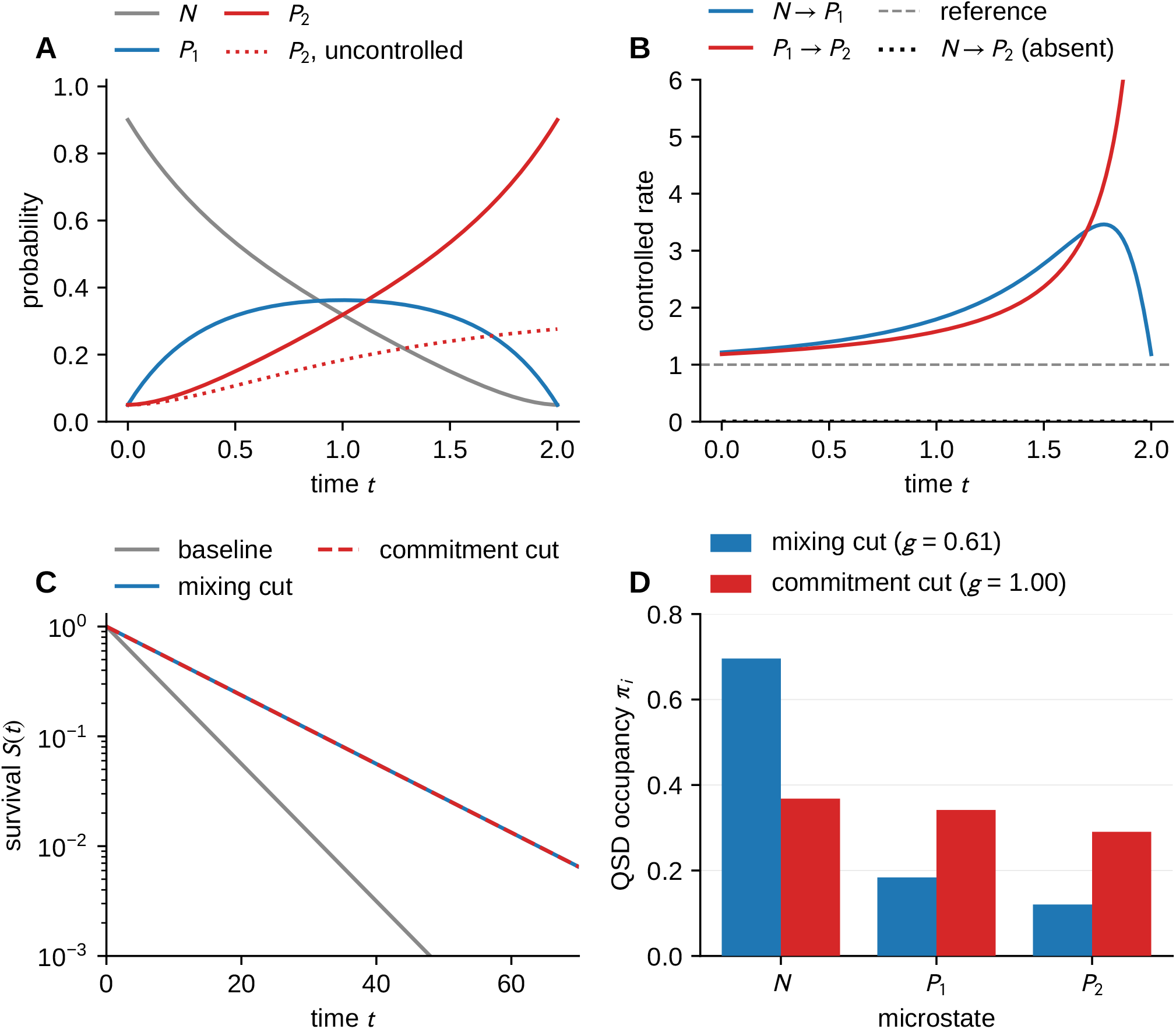
Minimum-effort steering and a mixing-versus-commitment cut. Three-state line *N ⇌ P*_1_ *⇌ P*_2_ *→ B* with all internal rates 1. (A) The Schrödinger bridge (*λ* = 0, horizon *T* = 2) drives the distribution *N → P*_1_ *→ P*_2_, whereas the uncontrolled process does not accumulate on *P*_2_. (B)The optimal control reweights only the existing edges, and the absent *N→ P*_2_ transition stays at zero, as the relative-entropy cost requires, so this panel checks the implementation rather than demonstrating a property of cells. (C) A mixing cut (reducing *α*) and a commitment cut (reducing *λ*) matched at Λ_*A→B*_ = 0.072 have coincident survival from the QSD. (D) They differ in occupancy (*π*_*P2*_ = 0.12 against 0.29) and in the gap of *Q*_*A*_ (0.61 against 1.00), since the mixing cut alters *Q*_*A*_.

**Figure S3.**
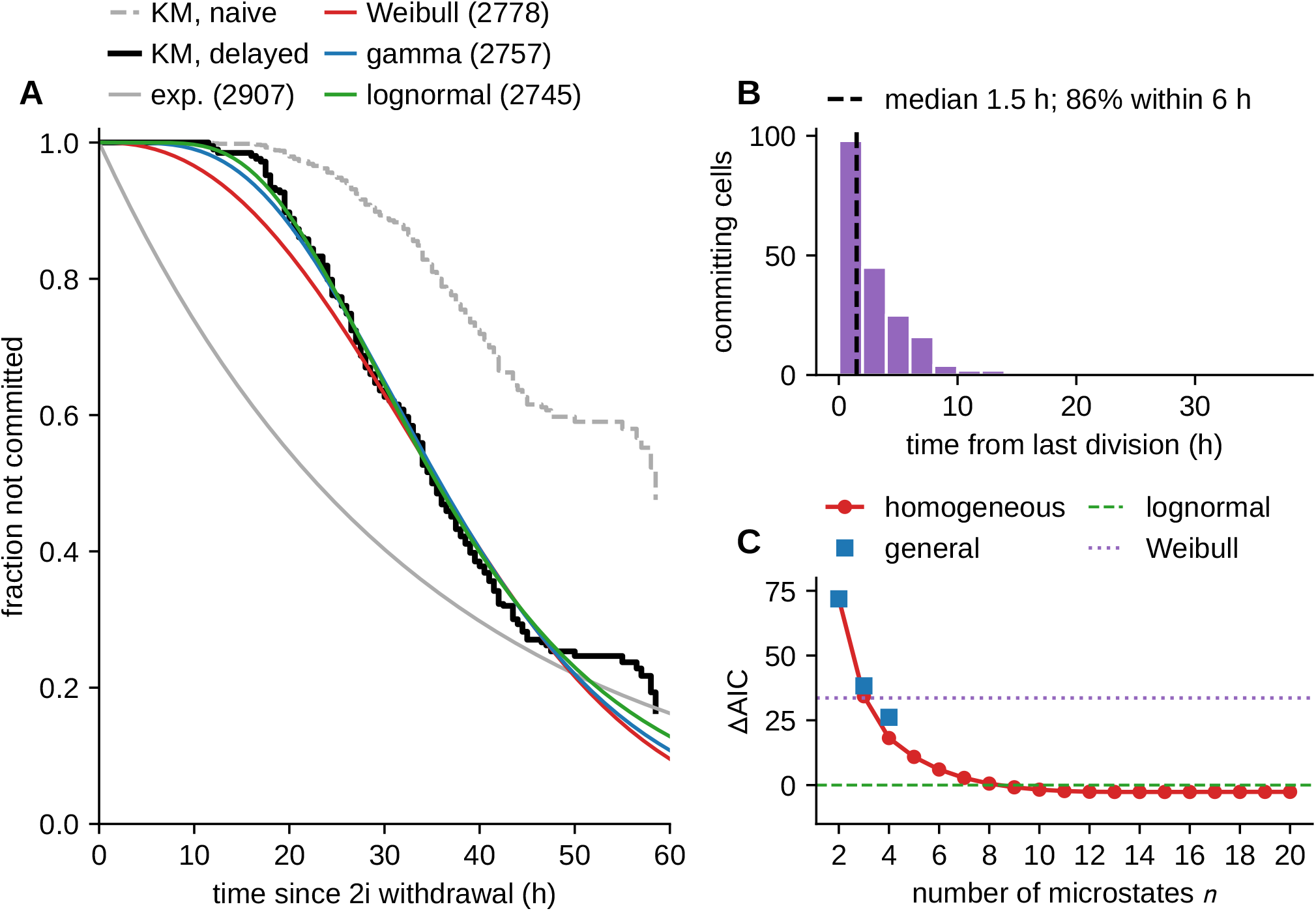
First-passage analysis of naïve exit. (A) Kaplan–Meier survival with every cell at risk from 2i withdrawal (dashed) and with each cell entered at its birth (delayed entry, solid), with lefttruncated parametric fits (AIC in parentheses). (B) Time from the last division to commitment, for all committing cells. (C) AIC of the killed line chains relative to the lognormal, against the number of microstates, computed with the interval-censored likelihood on the 30-min frame grid. Chains with free rates on every edge (squares) fit worse than homogeneous chains of the same length.

**Figure S4.**
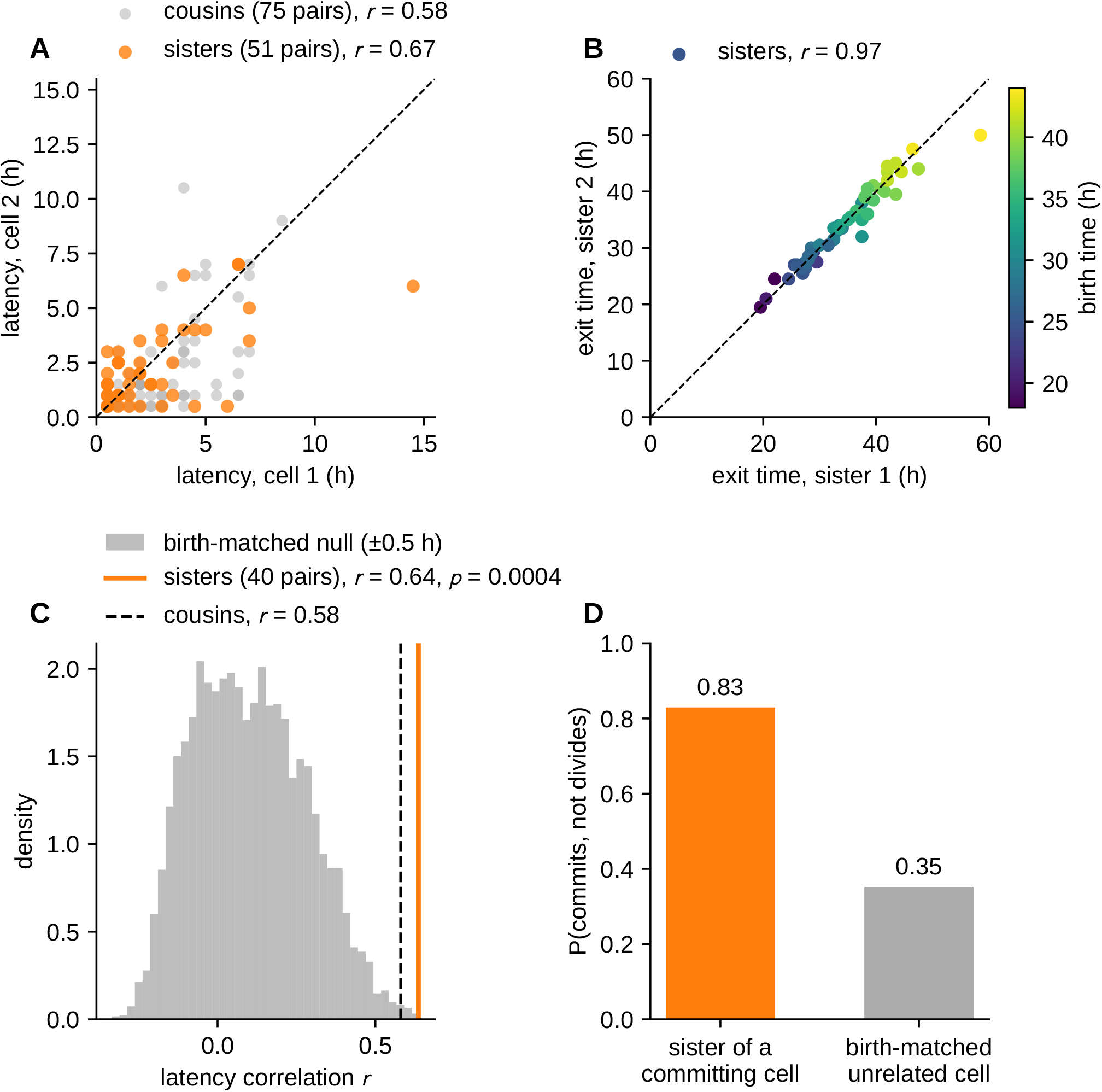
Correlation of commitment latency between relatives. (A) Latency of sisters and first cousins in which both commit. (B) Absolute exit times of sisters, coloured by birth time, whose near-perfect correlation reflects shared birth. (C) Correlation of sister latencies against the null distribution for unrelated committed cells matched on imaging date and birth time ( *±*0.5 h). (D) Probability that the sister of a committing cell commits rather than divides, against birth-matched unrelated cells.

**Figure S5.**
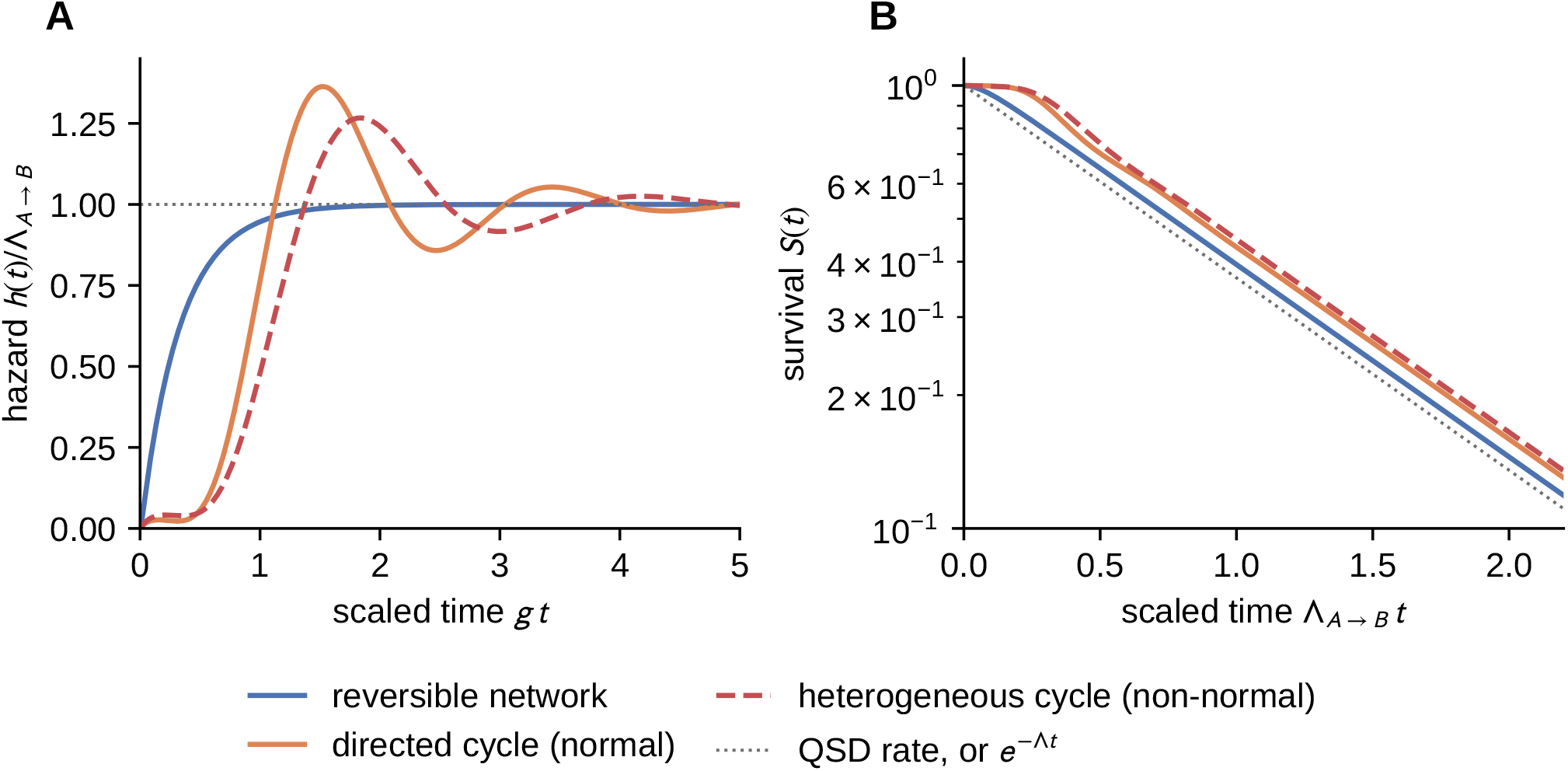
Departures from the reversible picture (Appendix D). The reversible elevenmicrostate network of Fig. 1B compared with a uniform directed cycle (normal, complex spectrum) and a heterogeneous directed cycle (genuinely non-normal), at matched stability index (*η* = 4.9, 4.2, 4.2) and started from a point mass far from the permissive set. (A) The exit hazard rises monotonically to Λ_*A*_*→*_*B*_ only in the reversible case. Both cycles overshoot the quasi-stationary rate (by 36% and 27% respectively) and ring down to it, so escape is transiently faster than the quasi-stationary reduction predicts. The uniform cycle is a circulant, hence normal, matrix, so the overshoot follows from its complex spectrum alone and requires no non-normality. (B) The corresponding survival curves depart visibly from the single exponential 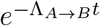 over the first mixing times, recovering it only asymptotically. The relative error in 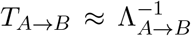 rises from 6.3% to 14.2% and 16.7%.

**Figure S6.**
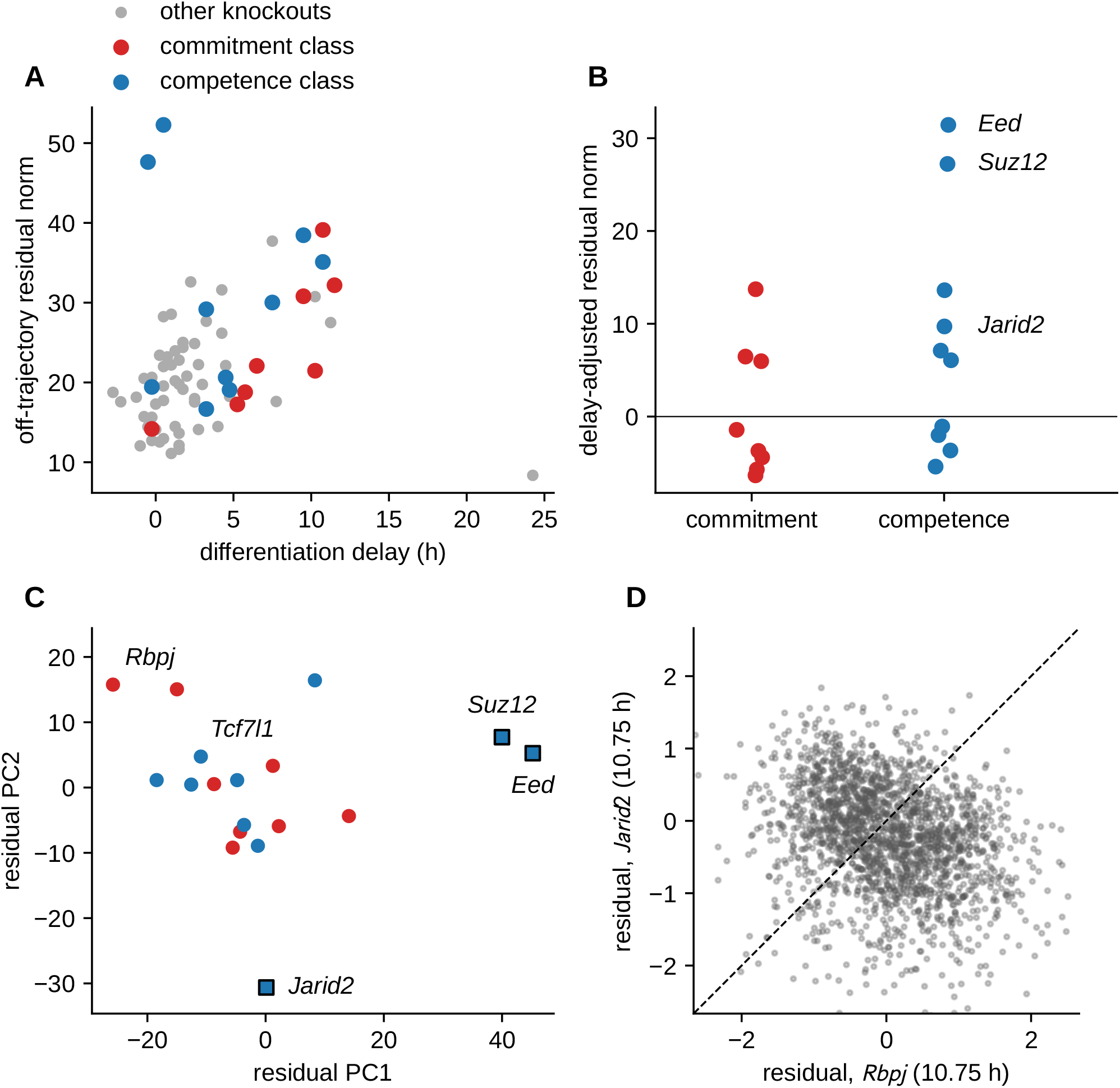
Equal differentiation delays do not fix internal state in the bulk knockout resource, but the channel is not resolved. Differentiation delay from differentially expressed genes, and the corresponding figure with the naïve-marker delay is produced by the same script. (A) Norm of each knockout’s off-trajectory residual against its delay. The delay accounts for 26% of signature variance. (B) Delay-adjusted residual norm by class (exact permutation *p* = 0.072, or 0.33 without PRC2). (C) Classified knockouts in the first two principal components of residual space. PRC2 lines are squares, and named knockouts are those discussed in the text (PERMANOVA *p* = 0.14, or 0.03 without PRC2). (D) Residuals of *Rbpj* and *Jarid2*, matched at a 10.75 h delay (*r* = *−*0.26).

